# Identifying HIV-1 RNA splice variant protein interactomes using HyPR-MS_SV_

**DOI:** 10.1101/2020.09.15.298190

**Authors:** Rachel A. Knoener, Edward L. Evans, Jordan T. Becker, Mark Scalf, Bayleigh E. Benner, Nathan M. Sherer, Lloyd M. Smith

**Author notes:** Senior Author (Nathan M. Sherer, Lloyd M. Smith).

## Abstract

HIV-1 generates unspliced (US), partially spliced (PS), and completely spliced (CS) classes of RNAs; each playing distinct roles in viral replication. Elucidating their host protein “interactomes” is crucial to understanding virus-host interplay. Here, we present HyPR-MS_SV_ for isolation of US, PS, and CS transcripts from a single population of infected CD4+ T-cells and mass spectrometric identification of their *in vivo* protein interactomes. Analysis revealed 212 proteins differentially associated with the unique RNA classes; including, preferential association of regulators of RNA stability with US- and PS-transcripts and, unexpectedly, mitochondria-linked proteins with US-transcripts. Remarkably, >80 of these factors screened by siRNA knock-down impacted HIV-1 gene expression. Fluorescence microscopy confirmed several to co-localize with HIV-1 US RNA and exhibit changes in abundance and/or localization over the course of infection. This study validates HyPR-MS_SV_ for discovery of viral splice variant protein interactomes and provides an unprecedented resource of factors and pathways likely important to HIV-1 replication.

## INTRODUCTION

The human immunodeficiency virus type 1 (HIV-1) uses the alternative splicing of a single primary RNA transcript to produce three major classes of viral RNA variants: unspliced (US), partially spliced (PS), and completely spliced (CS). The individual variants perform distinct roles during HIV-1 replication through dynamic interactions with specific viral and host proteins (Coffin et al., 1997). These protein “interactomes” guide the RNA through required cellular pathways encompassing splicing, RNA nuclear export, mRNA translation, and packaging of full-length, US RNA genomes into progeny virions that assemble at the plasma membrane. Each HIV-1 splice variant performs a distinct function and is thus predicted to interface with a unique protein interactome.

HIV gene expression is traditionally divided into two phases referred to as “early” and “late”. Early gene expression involves translation of auxiliary proteins Tat and Rev as well as the accessory protein Nef from CS transcripts. Tat and Rev localize to the nucleus where Tat facilitates viral transcription and Rev mediates nuclear export of intron-retaining US and PS transcripts. Late gene expression is marked by translation of the US transcript to synthesize Gag and Gag-Pol capsid proteins and translation of PS transcripts to generate Envelope glycoproteins as well as the Vpu, Vpr, and Vif immunomodulatory factors.

HIV splicing generates vast numbers of splice variants, with over 50 proposed to be physiologically significant (Emery et al., 2017; Ocwieja et al., 2012; Purcell and Martin, 1993; Vega et al., 2016). The locations of splice donor and acceptor sites (Sertznig et al., 2018; Vega et al., 2016), the identities of several *cis*- and *trans*-regulatory elements (Mahiet and Swanson, 2016; Sertznig et al., 2018; Stoltzfus, 2009), and the transcript and protein product abundances needed for efficient viral replication (Cullen, 1991; Karn and Stoltzfus, 2012; Weinberger et al., 2005) are still topics of intensive investigation toward the development of antiviral therapies.

Previous works have shown that the HIV splice variant classes interact differentially with both viral and host proteins. For example, HIV-1 US and PS transcripts hijack the cellular CRM1-mediated nuclear export pathway through the activities of Rev and a *cis*-acting RNA structure known as the Rev-response element (RRE) (Pollard and Malim, 1998). Rev multimerizes on the RRE and recruits CRM1 to form a functional RNA export complex (Bai et al., 2014; Daugherty et al., 2008; Daugherty et al., 2010; DiMattia et al., 2016; DiMattia et al., 2010; Fang et al., 2013). This is in contrast to CS transcripts that do not require Rev and recruit components of the NXF1/NXT1 export machinery, similar to the bulk of cellular fully-spliced mRNAs. A second example is HIV genome packaging wherein US transcripts are packaged into virions due to favored interactions between the Gag polyprotein and a structured RNA packaging signal known as “psi” in the 5’ untranslated region of the US transcript (Berkowitz et al., 1993; Lever et al., 1989; Luban and Goff, 1994). These binding sites are lost in PS and CS transcripts due to splicing (Purcell and Martin, 1993). Additional host factors are implicated as interactors in differential regulation of US, PS, and CS RNA transcripts (Bolinger and Boris-Lawrie, 2009; Freed and Mouland, 2006; Jin and Musier-Forsyth, 2019; Mbonye and Karn, 2014; McLaren et al., 2008; Meng and Lever, 2013; Swanson and Malim, 2006). However, the list is far from complete.

In the current study, we describe successful isolation of the three major HIV-1 splice variant classes from a single population of natively infected CD4+ T cells and define their distinct *in vivo* RNA-protein interactomes using mass spectrometry. We identify over 200 proteins differentially associated with the US, PS, and CS HIV splice variant pools, 116 of which are new virus-host interactors. Of these proteins, gene-specific siRNA knockdown indicated >80 new effectors of HIV-1 RNA regulation. We further demonstrate several instances of identified host protein co-localization with HIV RNA, and changes to the single-cell abundance and/or subcellular distribution of several identified host proteins over the course of HIV-1 infection. Collectively, we detail a powerful new approach for probing virus-host interactions and use it to expose many new host factors with potential roles in the HIV-1 replication cycle.

## RESULTS

### Purification of HIV-1 splice variant classes

We recently described HyPR-MS (**Hy**bridization **P**urification of **R**NA-Protein Complexes Followed by **M**ass **S**pectrometry); a strategy to identify the *in vivo* protein interactomes of specific viral RNAs, lncRNAs, and mRNAs (Knoener et al., 2017; Spiniello et al., 2018; Spiniello et al., 2019). Here, we present HyPR-MS_SV_, a strategy that expands the capabilities of HyPR-MS to differentiate *in vivo* protein interactomes for multiple splice variants (SV) derived from a single primary transcript and isolated from a single cell population. Applied here, we purified the three major classes of HIV-1 splice variants (unspliced (US), partially spliced (PS), and completely spliced (CS)) from a single population of infected Jurkat CD4+ T cells, then identified and characterized their protein interactomes using mass spectrometry.

To preserve *in vivo* viral RNA-protein complexes prior to cell lysis, Jurkat cultures were treated with formaldehyde at 48-hours post-infection (Multiplicity of Infection (MOI) of ~1 infectious units per cell). To isolate the US, PS, and CS RNA pools, three biotinylated capture oligonucleotides (CO’s) were designed complementary to three distinct regions of the HIV RNA genome: intron-1 (unique to US), intron-2 (present in both US and PS), and the 3’-exon (present in US, PS, and CS) (Figure 1A, Table S1). Cell lysates were first depleted of the US HIV RNA through hybridization to the intron-1 CO, followed by its capture with streptavidin-coated magnetic beads, and subsequent release using toehold-mediated oligonucleotide displacement. Additional hybridization, capture and release steps were subsequently repeated iteratively using first the intron-2 CO and then the 3’-exon CO for isolation of the PS and CS RNA pools, respectively. Once purified, proteins cross-linked to each isolated HIV RNA class were identified by mass spectrometry (Figure 1B).

**Figure 1:**
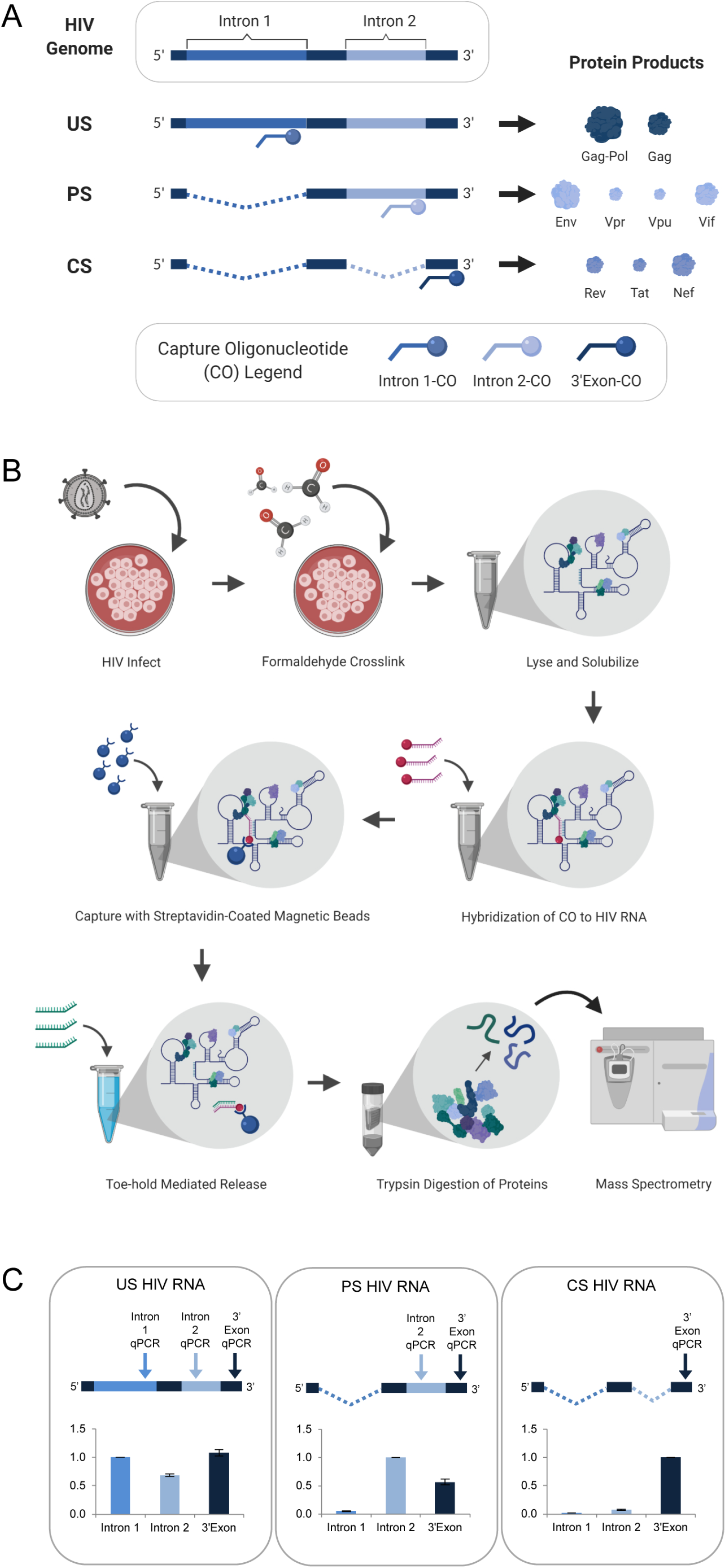
HyPR-MS for purification of HIV splice variant interactomes. **A.** COs were designed to complement specific regions of the HIV genome to make possible the isolation of the three HIV splice variant classes from one cell lysate. **B.** Overview of HyPR-MS_SV_ procedure. **C.** Purification of the HIV splice variant classes was verified using RT-qPCR assays specific to regions in Intron 1, Intron 2, and 3’Exon. The intensity data is normalized to the Intron 1 assay for US capture, the Intron 2 assay for PS capture and the 3’Exon assay for CS capture. Error bars are the standard deviation for three biological replicates.

RT-qPCR assays specific to intron 1, intron 2, and the 3’-exon (Table S1) were used to test RNA capture specificity and efficiency. For three biological replicates of the US, PS, and CS captures, the magnitude of amplification using each qPCR assay confirmed strong capture specificity for the desired splice variant class over the other two classes (Figure 1C, Table S2). Enrichment of HIV transcripts over a cellular control transcript (*GAPDH*) was >100-fold (Figure S1A, Table S2). Capture efficiency (the amount of each transcript depleted from the lysate after capture) was > 70% for each variant (Figure S1B, Table S2).

### Elucidation of unique protein interactomes for each HIV-1 splice variant class

To identify host proteins differentially interacting with the US, PS, and CS RNA pools, we isolated the *in vivo* crosslinked HIV RNA variants from three biological replicate experiments of 5×10^7^ infected Jurkat cells; with each replicate generated from a separate set of cultured cells and virus preparations. Interacting proteins from the US, PS, and CS capture samples were purified, analyzed by bottom-up mass spectrometry, then identified and quantified using search and label-free quantitation algorithms (Cox and Mann, 2008; Tyanova et al., 2016). We determined which proteins preferentially associated with each splice variant class by conducting three pairwise comparisons: US vs PS, US vs CS, and PS vs CS. Using the Student’s t-test and a permutation-based false discovery rate (FDR) of 5%, we identified 212 proteins that differentially interacted with one or more of the HIV splice variant classes; 101, 93, and 68 proteins in the US, PS, and CS captures, respectively (Table S3). Hierarchical clustering was used to organize the 212 proteins into a heat map for visualization. The associated dendrogram shows the extent of similarity among the “interaction profiles” for each protein. This analysis revealed clusters of proteins elevated for each individual class as well as proteins common to members of the three HIV splice variant classes. The most abundant of these were proteins preferentially associated with both the US and PS HIV transcripts but not the CS pool (45 proteins) (Figure 2A).

**Figure 2:**
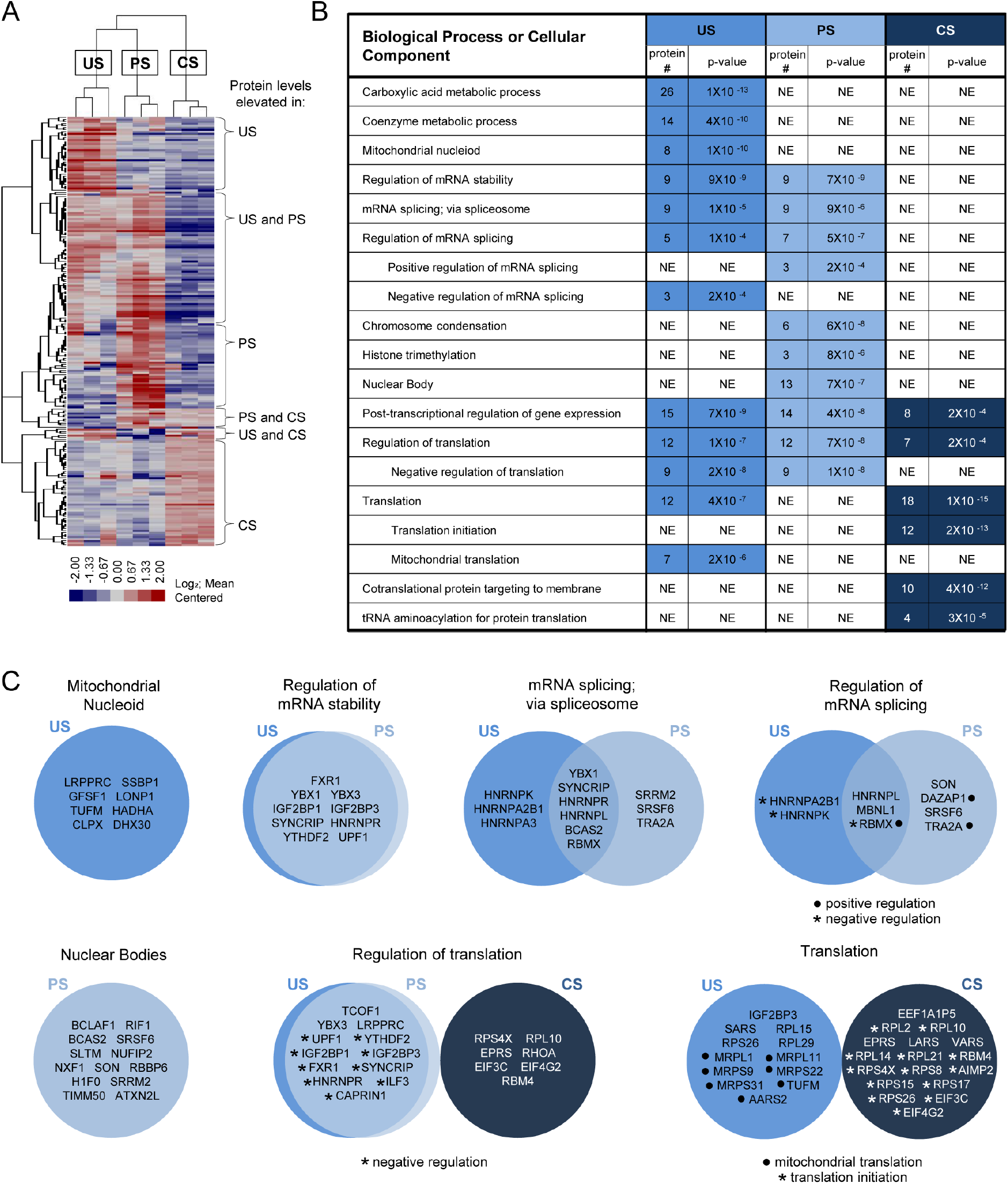
Determination and analysis of HIV splice variant protein interactomes. **A.** Heat map depicts relative intensities for each of the 212 proteins (rows) in each of the three biological replicates of the US, PS, and CS (columns) differential interactomes. **B.** Condensed list of gene ontology biological process or cellular component terms enriched in each of the HIV splice variant interactomes. The “protein #” column indicates the number of proteins in the interactome that are annotated with the biological process indicated. The “p-value” column indicates the likelihood that the proteins of the biological process are present in each interactome by random chance and were provided by GO term enrichment software (Mi et al., 2017). A lower p-value suggests non-random over-representation of a biological process.”NE”= Not Enriched. **C.** Venn Diagrams of proteins annotated for biological processes or cellular components enriched in the splice variant differential interactomes.

We used these interactome data to infer biological pathways potentially relevant to the regulation of each splice variant class. Using gene ontology (GO) term enrichment algorithms (Mi et al., 2017), we evaluated each interactome for enrichment of proteins involved in specific biological processes. This analysis revealed over-representation of several GO terms in the interactome of each splice variant class; some common to more than one class (Table S4, Figure 2B). Notable amongst these were nine proteins known to regulate RNA stability, associated with both the US and PS, but not CS, transcripts (FXR1, YBX1, YBX3, IGF2BP1, IGF2BP3, SYNCRIP, HNRNPR, YTHDF2, and UPF1). Proteins involved in mRNA splicing, and the regulation thereof, were also elevated in the US and PS relative to the CS capture samples (YBX1, SYNCRIP, HNRNPR, HNRNPL, BCAS2, RBMX, and MBNL1) but with less congruence. A subset of these proteins were elevated only in the US capture samples (HNRNPA2B1 and HNRNPK; negative regulators of splicing) or PS capture samples (DAZAP1 and TRA2A; positive regulators of splicing), but not in both. PS captures were also exclusively enriched for proteins found in nuclear bodies. For cytoplasmic activities, proteins involved in translation were highly enriched in both the US (12 proteins) and CS (18 proteins) interactomes. Interestingly, however, while the CS interactome included translation initiation proteins (as may be expected), the US interactome was enriched for proteins linked to mRNA translation in the mitochondria, with cellular component GO term enrichment analysis further revealing 45 mitochondrion proteins enriched in the US RNA interactome, 8 of which are mitochondrial nucleoid proteins (Figure 2C, Table S4).

### Validation of HyPR-MS_SV_ defined HIV-1 RNA interactors using RNA silencing

To determine their potential relevance to HIV-1 gene expression, 121 host proteins identified by HyPR-MS_SV_ were targeted for siRNA knockdown (KD; Table S5) in HEK293T cells and passed cell viability criteria (Table S6). Following KD, cells were infected with a 2-color HIV-1 virus engineered to report single-cell levels of viral US (Gag-CFP) and CS (mCherry) gene expression (Figures 3A and 3B) (Knoener et al., 2017). Relative to a scrambled siRNA control, statistically significant changes (p-value <0.05) to early (CS) and/or late (US) gene expression were observed for a remarkable 69% (84 total) of the targeted host genes (Figure 3C; Table S7). The KD of 33 host proteins affected the expression of US and CS protein products in the same direction (either both increased or both decreased) and with approximately the same magnitude. By comparing mCherry:CFP fluorescence ratios for each protein KD to the negative control, we determined that CS and US protein expression were differentially affected by KD of 51 host proteins; for 26 of the proteins the expression changes were in the same direction but with different magnitudes; for 18 only the expression of the US RNA protein product was affected; and for 7 only the expression of the CS RNA protein product was affected (Figure 3C, Table S7). Based on the direction of the changes in HIV-1 gene expression (increased or decreased), we categorized 71 host genes as putative “negative” effectors and 15 as putative “positive” effectors (Figure 3C, Table S7). Interestingly, of the 16 negative effectors,10 were implicated in mitochondria-associated pathways based on GO analysis; of those ten, nine were identified by HyPR-MS_SV_ to preferentially interact with the US HIV RNA (Tables S4 and S7).

**Figure 3:**
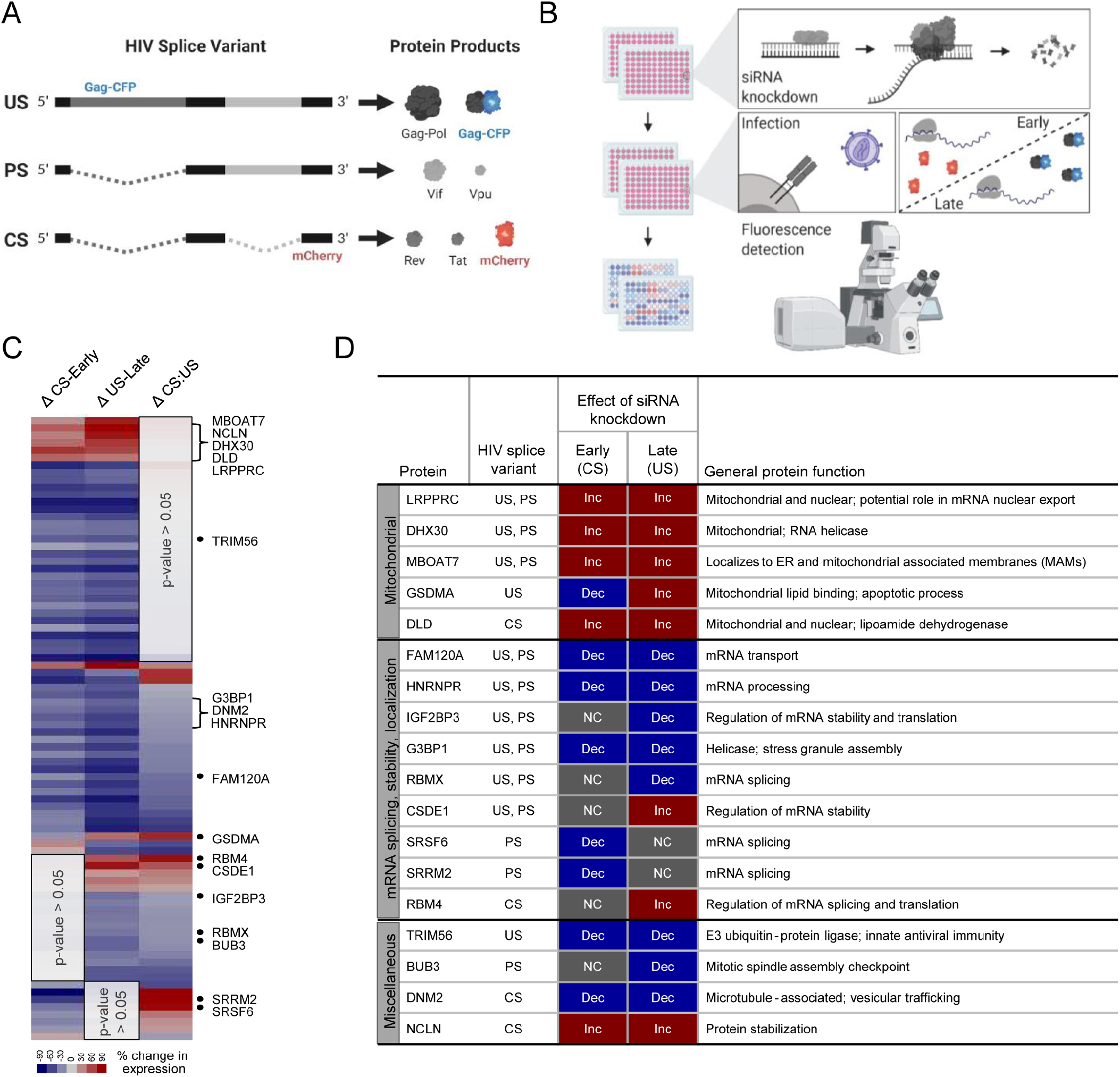
Screen for host protein effects on early and late HIV gene expression. **A.** The HIV-1 reporter virus used contains a Gag open-reading frame (ORF) with three copies, in tandem, of a cyan fluorescent protein (CFP) reporter and an mCherry reporter in the Nef ORF. **B.** In 96-well plates, 293T-ACT-YFP cells were transfected with gene-specific siRNAs for 4 hours, 48-hours later they were transfected again for 4 hours followed by incubation with the HIV reporter virus. The cells were then fixed at 48-hours post incubation. Fluorescence microscopy was used to quantify CFP and mCherry. **C.** Heatmap of HIV gene expression changes after siRNA knockdown of host proteins. Eighty-four of 121 proteins showed statistically significant changes in early and/or late HIV gene expression (p-value <0.05). **D.** Twenty proteins were selected for confirmation of KD efficacy using western blot. The table summarizes the HyPR-MS and KD results for the proteins for which the WB or IF showed significant KD of the targeted host protein (18 proteins). Note: KDs detection for proteins IGF2BP3, SRRM2, and DNM2 were unsuccessful by WB but were later shown to be effective using the same antibodies in fixed cell immunofluorescence (Tables S9 and S11).

### HyPR-MS_SV_ candidates co-localize with US HIV RNA at distinct subcellular locations

We selected a subset of 20 HyPR-MS_SV_ identified host proteins for further validation studies. This subset was, in part, chosen based on the commercial availability of antibodies that allowed for immunoblot- and/or immunofluorescence-based detection of the host proteins (Table S8), and included five proteins linked to mitochondria (LRPPRC, DHX30, MBOAT7, GSDMA, and DLD; all negative effectors of US RNA gene expression); ten genes encoding proteins with functions related to mRNA processing, localization and stability (FAM120A, HNRNPR, IGF2BP3, G3BP1, RBMX, CSDE1, SRSF6, SRRM2, RBM4, and RPL15; the majority of which were positive effectors of either US or CS gene expression); and five additional proteins that had not previously been linked to RNA regulation (TRIM56, BUB3, DNM2, DYNC1H1 and NCLN) (Figure 3D, Table S7). Fifteen of the 20 proteins were detected by immunoblot and siRNA KD was confirmed (31 to 95% relative to negative control siRNA) (Figure S2). IGF2BP3, SRRM2, DNM2, RPL15, and DYNC1H1 KDs were not confirmed by immunoblot (Tables S9).

US HIV RNA-protein interactions may commence as early as production of the nascent HIV transcript in the nucleus or as late as virus particle formation at the plasma membrane. To determine potential sites of interaction, we used 3-color combined fluorescence in situ hybridization / immunofluorescence (FISH/IF) single-cell imaging to show host factor subcellular localization relative to US RNA and viral Gag proteins (Figures 4A and S3). Cells were infected with an HIV-1 reporter virus (HIV-1 E-R-CFP) allowing for identification of infected cells and confirmation of specificity of the US RNA FISH probes (Stellaris FISH probe set specific to intron-1 (Table S10)) and Gag antibody (anti-p24Gag (Table S8)). Host proteins were detected using the primary antibodies employed for our immunoblot analysis (Table S8); with 17 of the 18 host proteins (all but DLD) detected by IF and showing greater than 40% decreases in IF signal after host protein siRNA KD. This imaging-based analysis also allowed verification of the efficacy of siRNA KD for three of the host proteins (IGF2BP3, SRRM2, and DNM2) that we had been unable to detect using immunoblot (Table S11).

**Figure 4:**
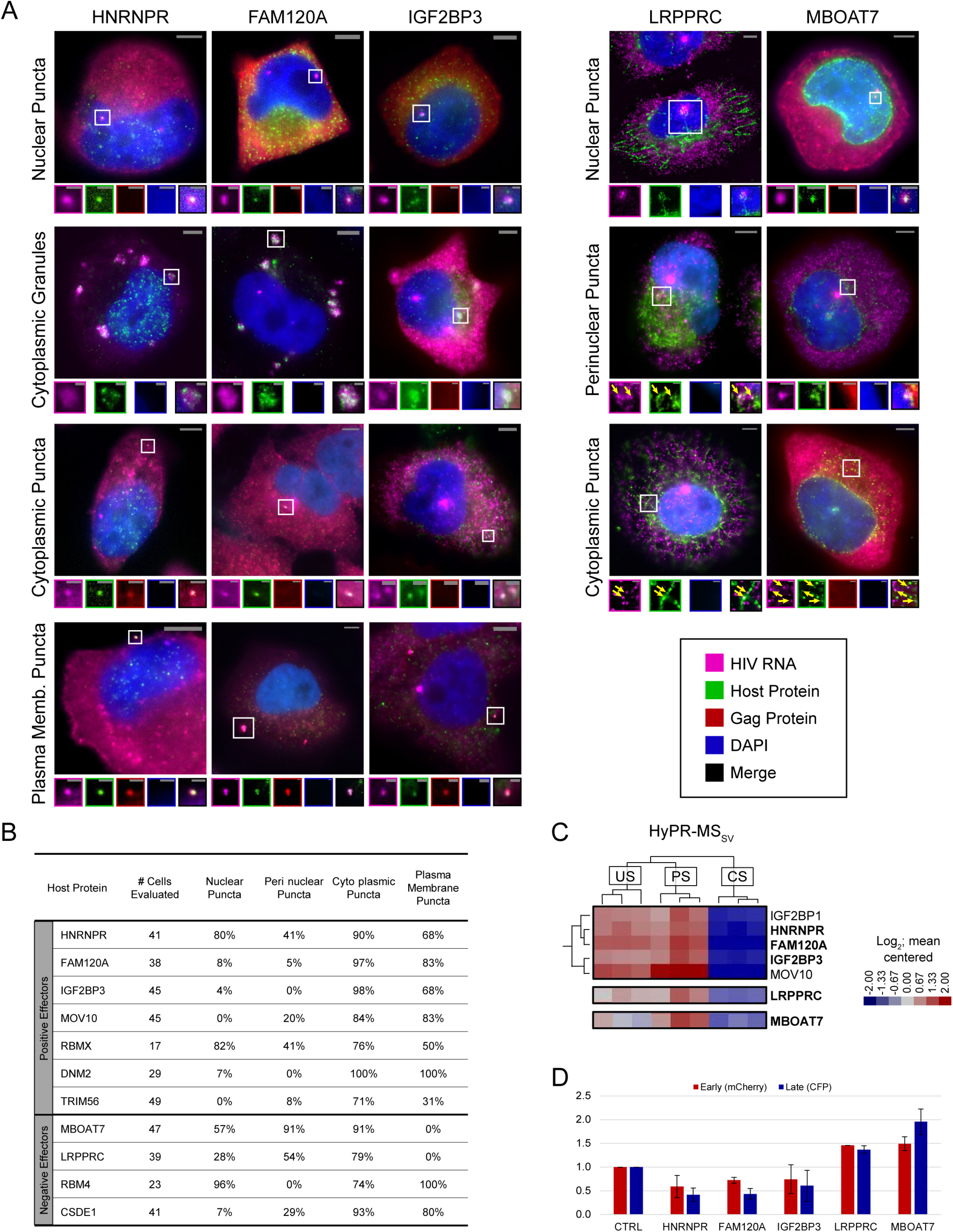
US HIV RNA co-localizes with positive and negative effectors at multiple sites within the cell. **A.** Representative images of co-localization phenotypes observed using FISH/IF. For each, a merged image of a cell highlighting a site of co-localization (white square) is shown. Enlarged regions of interest (ROI) of each fluorescence channel are displayed in the associated small panels to separate overlapping US HIV RNA, host protein, HIV Gag polyprotein, and DAPI signals. Some images were obtained from experimental replicates that did not include Gag IF and therefore do not include images from the corresponding channel. Note: Brightness and contrast settings were adjusted individually for each color channel of the images to effectively show co-localization. These settings may be different for the ROIs. **B.** Table showing the frequency of observing a particular co-localization phenotype of US HIV RNA with each of 11 host proteins. Frequencies are displayed as the percentage of cells observed with each co-localization phenotype. **C.** Regions-of-interest from the HyPR-MS, hierarchically clustered heatmap (Fig. 2A) showing the close relation of HNRNPR, FAM120A and IGF2BP3 interaction profiles; each preferentially interacted with US and PS HIV RNA. **D.** Data for proteins of interest from the siRNA KD screen. HIV gene expression decreases for HNRNPR, FAM120A, and IGF2BP3 upon KD with a greater decrease in late gene expression than in early. For mitochondria-related proteins LRPPRC and MBOAT7, HIV gene expression increases upon KD of the host protein.

FISH/IF was performed on HeLa cells 48-hours post-infection to localize US RNA, Gag, and each of the 17 HyPR-MS_SV_ identified host proteins. Analysis by single-cell fluorescence microscopy showed consistent co-localization of US HIV-1 RNA with 11 of the proteins (Figures 4A and S3), with four (HNRNPR, RBMX, RBM4, MBOAT7) predominantly localized to the nucleus or near the nuclear membrane and seven (FAM120A, IGF2BP3, MOV10, TRIM56, DNM2, LRPPRC, CSDE1) predominantly localized to the cytoplasm in uninfected cells (Figure S4). In infected cells, we observed five recurrent US RNA-host protein co-localization phenotypes: (1) at nuclear puncta; (2) at puncta proximal to the nuclear membrane; (3) at cytoplasmic puncta; (4) at large, cytoplasmic complexes reminiscent of stress granules, and; (5) at the plasma membrane (Figures 4A and S3). In the nucleus, we typically observed one or two bright US RNA puncta per cell, consistent with prior reports describing sites of active HIV-1 transcription (Puray-Chavez et al., 2017). Puncta proximal to the nuclear membrane and cytoplasmic puncta were smaller, more numerous, and of lower intensity. Cytoplasmic granules were large with moderate intensity accumulations of US HIV RNA surrounded by or spotted with host protein. Plasma membrane puncta were variable in size and intensity and often co-localized with Gag, thus likely represent virion assembly sites (Figures 4A and S3).

We quantified the frequency of each co-localization phenotype for 17-52 cells per antibody (Figure 4B, Table S12), excluding cytoplasmic granules that were only rarely observed. The data revealed that proteins that predominantly localize to the nucleus or proximal to the nuclear membrane (HNRNPR, RBMX, RBM4, MBOAT7) had a higher frequency of co-localization with HIV RNA at nuclear puncta (57-96%) relative to proteins that were predominantly localized to the cytoplasm (FAM120A, IGF2BP3, MOV10, DNM2, TRIM56, LRPPRC, CSDE1; 0-8%). Two proteins (HNRNPR and RBMX) co-localized frequently with HIV-1 US RNA at all four quantified sites (41-90%). All 11 host proteins co-localized with US RNA at small cytoplasmic puncta in a high percentage of cells (71-100%) with most (all but LRPPRC and MBOAT7) co-localizing with US RNA at the plasma membrane (31-100% of cells), generally with Gag also present (Figure 4B, Table S12).

### A subset of HyPR-MS_SV_ candidates likely co-traffic with US RNAs from sites of transcription to the cytoplasm

Several HyPR-MS candidates (HNRNPR, FAM120A, IGF2BP3, RBMX, RBM4, CSDE1, DNM2, LRPPRC, MBOAT7) were observed to accumulate at bright US RNA nuclear puncta, suggesting that they associate with US RNA at or near sites of *de novo* transcription (Figure S3). Of these, HNRNPR, FAM120A, and IGF2BP3 were of particular interest because all three exhibited four US HIV RNA co-localization phenotypes (nuclear puncta, cytoplasmic granules, cytoplasmic puncta, and plasma membrane puncta) (Figure 4B); preferentially interacted with US and PS, but not CS, HIV RNA as determined by HyPR-MS_SV_ (Figure 4C, Table S7), and positively affected US but not CS gene expression upon siRNA KD (Figure 4D, Tables S7). By contrast, LRPPRC, a protein shown to localize to the mitochondria as well as the nucleus (Mili and Pinol-Roma, 2003; Ruzzenente et al., 2012), and MBOAT7, a protein shown to localize to mitochondria-associated membranes (Hirata et al., 2013), localized to US HIV RNA nuclear puncta, at perinuclear puncta, and at cytoplasmic puncta but were not observed to co-localize with US RNA and Gag at the plasma membrane. Similar to the HNRNPR, FAM120A, and IGF2BP3, both LRPPRC and MBOAT7 were preferentially associated with US and PS relative to CS transcripts based on HyPR-MS_SV_ analysis. However, unlike HNRNPR / FAM120A / IGF2BP3, each of these proteins were negative effectors of both US and CS HIV-1 gene expression (Figure 4D, Table S7). Interestingly, LRPPRC was detected not only near transcription sites but also in a trail-like pattern that extended to the periphery of the nucleus (Figures 4A and S3) and MBOAT7 was observed at transcription sites, at smaller subnuclear US HIV RNA puncta, and with high frequency and abundance at US RNA puncta at or near the nuclear membrane (Figures 4A and S3).

### HIV-1 infection alters the abundance and localization of several HyPR-MS_SV_ identified proteins

The FISH/IF single cell analyses of US HIV RNA, Gag, and host proteins also allowed for tracking of host factor responses to infection (Figure 5). For example, HNRNPR, generally a nuclear protein, was primarily localized to the nucleus of cells expressing no, or low amounts of, Gag and US RNA, but exhibited marked shifts from the nucleus to the cytoplasm in cells with high levels of Gag and US RNA expression (Figure 5A). Changes to MBOAT7 were also striking, with much higher levels of expression in cells with abundant Gag and US RNA than in uninfected or early infected cells (Figure 5B).

**Figure 5:**
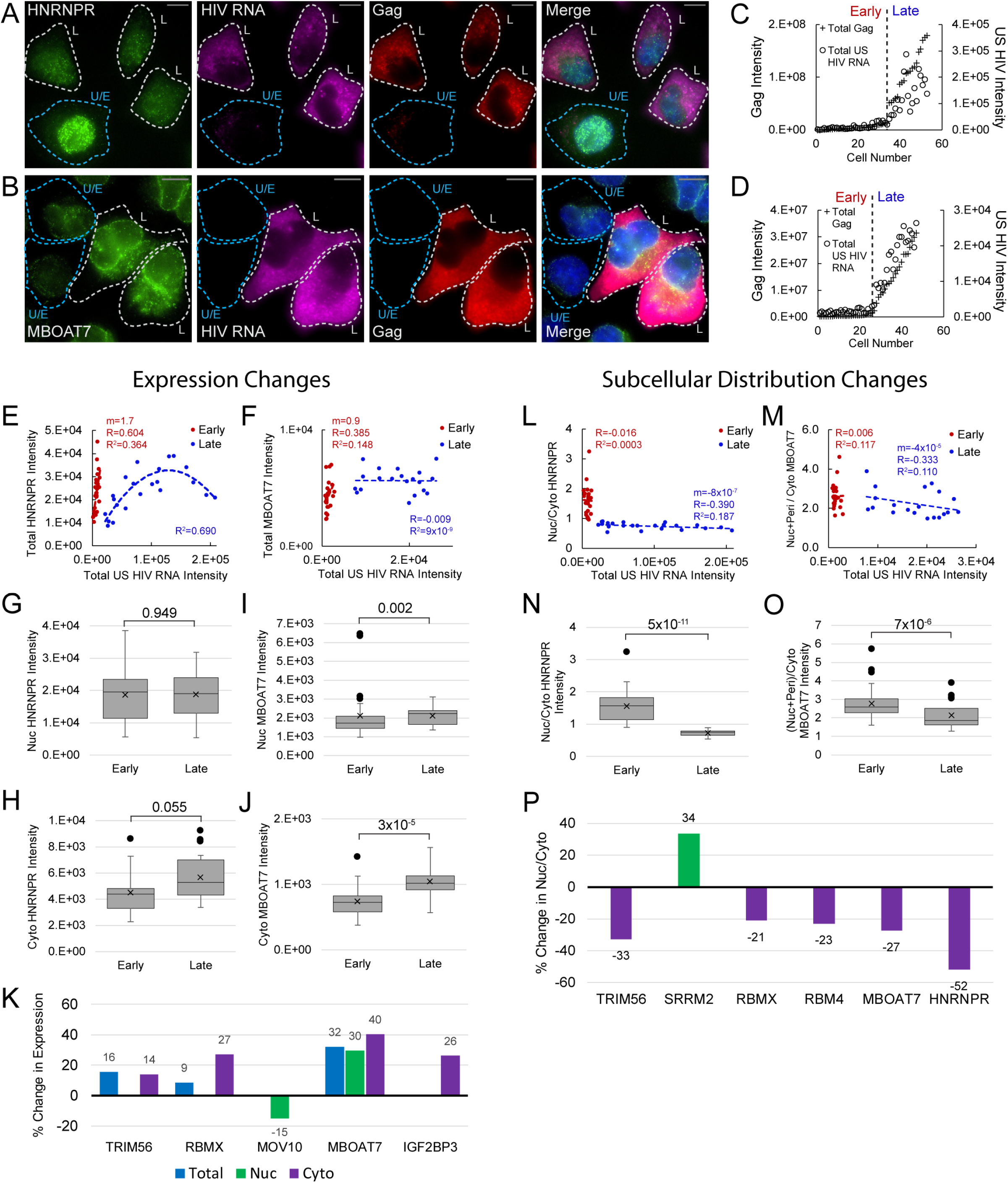
Host protein expression and cellular distribution. **A.** HNRNPR cellular distribution appears to be different in uninfected/early stage infected cells (U/E; blue outlines) than in late stage infected cells (L; white outlines). **B.** MBOAT7 expression appears greater in late stage infected cells than in uninfected/early stage infected cells. **C-D.** Plots of cellular Gag-IF and US HIV RNA-FISH intensities for cells analyzed for HNRNPR (C) and MBOAT7 (D). Cells prior to the inflection points in each plot are termed “early cells” as they are either uninfected or at stages of HIV replication prior to late gene expression (i.e. high amounts of Gag in the cytoplasm). Cells after the inflection point are termed “late cells” as they express high, IF-detectable levels of Gag in the cytoplasm. **E-F.** For each cell in the early and late cell sub-groups, the intensity of US HIV RNA versus the intensity of HNRNPR (E) and MBOAT7 (F) is plotted. Linear or polynomial regressions (R^2^) are fit to each early and late sub-group and the Pearson’s correlation coefficient (R) calculated for linear regressions. This demonstrates the extent of correlation between US HIV RNA expression and the expression of each host protein. **G-J.** A student’s t-test is applied to determine if the host protein intensities in early cells are significantly different from those in late cells in the nucleus and the cytoplasm. **K.** % change in the median expression for all host proteins with early vs late p-values < 0.05. Calculations were made for total cell, nuclear, and cytoplasmic differences. **L-M.** Total cellular US HIV RNA intensities versus host protein nuc/cyto or nuc+peri/cyto ratios. Demonstrates extent of correlation of host protein cellular distribution with US HIV RNA expression. **N-O.** A student’s t-test measures significant differences in the cellular distribution between the early and late cells for HNRNPR and MBOAT7. **P.** % change in the median nuc/cyto or nuc+peri/cyto ratio for all host proteins with p-values < 0.05. Purple indicates the late cells have a higher proportion of the host protein in the cytoplasm than do the early cells. Green indicates the late cells have a higher proportion in the nucleus.

To further track HyPR-MS_SV_ host factor changes, we plotted single-cell measurements of total Gag and total US HIV RNA and used the resulting inflection point to discriminate cells in “early” and “late” stages of HIV gene expression (Figures 5C and 5D; Table S13). We measured relative host protein abundances for twelve of these factors at these stages (HNRNPR, FAM120A, IGRF2BP3, LRPPRC, MBOAT7, CSDE1, DNM2, MOV10, RBM4, RBMX, SRRM2, TRIM56 (Figure 5E and 5F, Table S13)). In general, each host protein exhibited non-random, bimodal expression changes from “early” and “late” HIV gene expression (Figure S5, Table S13). For example, in early/uninfected cells we observed linear increases in HNRNPR and MBOAT7 expression, positively correlating with the subtle increases in US HIV RNA expression (Figure 5E and 5F; slope m=1.7, 0.9 respectively). However, in late cells, HNRNPR expression rose then fell again as per-cell US RNA increased, fitting a polynomial rather than linear, trendline (R^2^=0.690) (Figure 5E) and suggesting changes to cell signaling; while MBOAT7 expression levels plateaued (Figure 5F, slope m=-0.007).

A similar analysis was performed after image-based segmentation of cells into nuclear and cytoplasmic compartments to better discriminate the subcellular location in which host protein changes occurred (Figure S5, Table S13). For HNRNPR, the same trends were observed in the nucleus and cytoplasm as were seen for the total cell (Figure S5). For MBOAT7, nuclear expression plateaued as it did for total cell expression, but the cytoplasmic expression increased slightly as US RNA and Gag abundance increased (Figure S5). In all, the expression of each of the 12 host proteins showed significant correlation with the expression of US HIV RNA in at least one of the following sub-groups: early-nuclear, late-nuclear, early-cytoplasmic, late-cytoplasmic (Figure S6, Table S13).

For HNRNPR and MBOAT7, the most evident differences were in cytoplasmic expression (cyto HNRNPR, median increase=21%, p=0.055; cyto MBOAT7, median increase=41%, p=3×10^-5^) (Figure 5G–5J, Figure S5, Table S13). In all, changes to nuclear or cytoplasmic abundance were observed for five host proteins (p-values < 0.05; MBOAT7, TRIM56, RBMX, MOV10, and IGF2BP3) (Figures 5K, S5, Table S13). Three of these proteins showed differences to total cellular expression (MBOAT7, RBMX, and TRIM56), with RBMX and TRIM56 only increasing in the cytoplasm. Two proteins did not show net differences in overall expression, but exhibited statistically significant differences (p-value < 0.05) in expression in the nucleus (MOV10) or the cytoplasm (IGF2BP3).

To identify potential host protein translocation events, we evaluated single cell nuclear-to-cytoplasmic (nuc/cyto) ratios relative to US RNA abundance and looked for statistically significant differences in early and late cells (Figure S5, Table S13). HNRNPR nuc/cyto ratios ranged from 1 to 3 in early/uninfected cells but only ranged from 0.6 to 0.9 in late infected cells; exhibiting a negative correlation with US HIV RNA expression (Figure 5L). For MBOAT7, the nuc/cyto ratio ranged from 1.7 to 3.9 in early cells and 1.5 to 3.9 in late cells; with no significant correlation with US RNA expression for either phase (Figure 5M). However, overall nuc/cyto ratios were significantly lower for late cells relative to early cells for both proteins (median decrease=-52%; p=5X10^-11^ and median decrease=-27%; p=7X10^-6^, respectively) (Figure 5N and 5O). In all, the nuc/cyto ratios of six HyPR-MS candidate proteins showed notable changes to nuc/cyto ratio (HNRNPR, MBOAT7, TRIM56, SRRM2, RBMX, and RBM4); all, with the exception of SRRM2, exhibiting relative increases to cytoplasmic abundance (Figure 5P, Table S13).

Taken together, these analyses demonstrated that many of the host factors identified by HyPR-MS_SV_ not only modulate HIV-1 gene expression (Figure 3) but co-localize with HIV-1 US RNA (Figure 4) and respond to infection by increasing in abundance and/or undergoing alterations to subcellular distribution (Figure 5).

## DISCUSSION

The variation in gene products encoded by the HIV-1 genome is largely achieved through regulated synthesis of a diverse RNA transcriptome. Deciphering the distinct cellular processes each splice variant undergoes and the host proteins involved is critical to understanding HIV-1 replication. Here, using HIV-1 as a relevant model system, we describe HyPR-MS_SV_ as a new tool that can be applied to elucidate distinct protein interactomes for distinct splice variant classes.

Isolation of the multiple HIV splice variant classes and comparative analysis of their differential protein interactors yielded a rich interactome resource valuable for studies of HIV-1 RNA metabolism. Notably, the protein interactomes of the three splice variant classes differ markedly from one another, presumably reflecting functional differerences (Figure 2). We uncovered over fifty proteins that differentially impacted early and late HIV gene expression based on siRNA KD (Figure 3), mapped the cellular locations where several of the host proteins co-localized with US HIV RNA (Figure 4), and established a correlation between infection and altered levels of expression or subcellular localization for several host proteins (Figure 5). Combined, these results provide a road-map for RNA-protein interactions potentially central to HIV replication (Figure 6).

**Figure 6:**
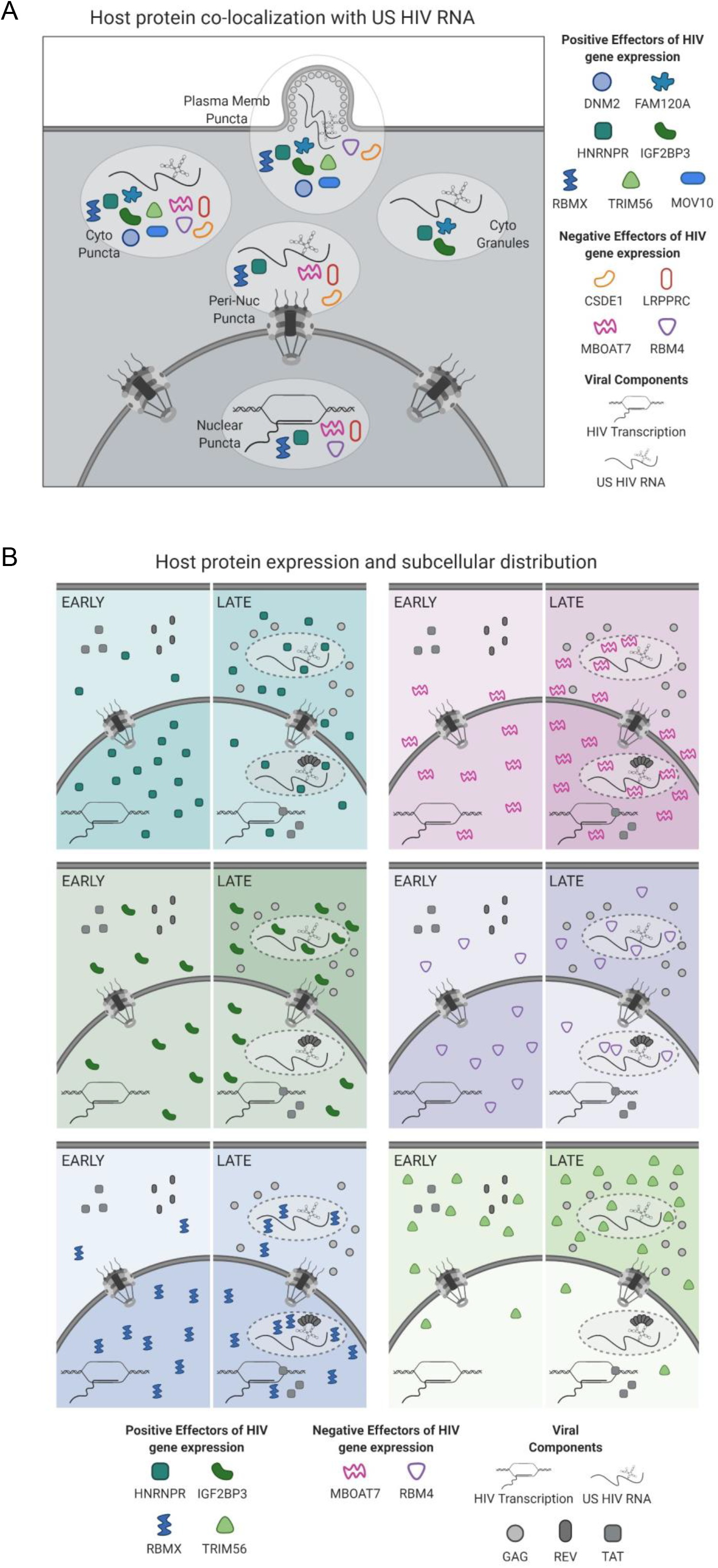
Models for subcellular co-localization, expression, and distribution of host proteins. **A.** Summary of cellular co-localization phenotypes observed by FISH/IF analysis of HIV US RNA and select host proteins. Host proteins represented include both positive and negative effectors of HIV gene expression as was determined by siRNA knockdown. **B.** Models of host protein changes in expression and cellular distribution from early to late HIV gene expression. Host protein quantities represented here show the general, but not exact, scale of changes in host protein abundance in the nucleus and cytoplasm over the course of HIV replication. The extent of shading in the nucleus or cytoplasm correlates with that same change in abundance.

In developing the HyPR-MS_SV_ approach, we aimed to (1) ensure the relevance of protein interactors by only pursuing interactions that occur in cells (i.e., *in vivo*) (2) ensure versatility of the technique for broad applications wherein it is useful to differentiate between one or more RNA splice variants for comparative RNA-capture proteomics and (3) determine, for the first time, the protein interactomes for the three major HIV splice variant classes. The first two goals were achieved by configuring HyPR-MS_SV_ to sequentially deplete specific classes of HIV RNA from the same pool of natively infected cell lysates using three independently-targeted sets of short (~30 nt) biotinylated capture oligos (Figure 1). To our knowledge, all previous studies for discovery of HIV RNA protein interactors, with the exception of our prior study (Knoener et al., 2017), utilized synthetic viral RNAs as bait added to cellular lysates (Marchand et al., 2011; Singh et al., 2016) or viral constructs engineered to encode artificial RNA sequences for the purpose of RNA “tagging” (*e.g*., MS2 loops) (Kula et al., 2011). While effective at identifying protein interactors, both of these strategies may complicate interpretation of results either by eliminating the cellular context of interactions (*e.g*., some interactions occur in the nucleus and some in the cytoplasm) or by introducing non-native components that can interfere with native interactions. Additional strengths of the HyPR-MS_SV_ approach are that it can be used to extract native RNA transcripts produced from any strain or infected cell type and can easily be adapted to study other viruses or cellular RNA splice variants.

By isolating the HIV-1 splice variant classes, we were able to compare RNAs with both shared and distinct sequences, and likely corresponding secondary, tertiary, and quaternary structures, to decipher how their protein interactors may consequently differ. We validated the sequential capture of US, PS, and CS HIV RNAs using RT-qPCR and showed at least 200-fold specificity relative to cellular RNAs and at least 10-fold specificity for the splice variant class of interest in each capture. We determined using statistical analysis that over 200 proteins interact preferentially with any one subset of the HIV splice variant classes at 48 h post-infection. Among these are proteins specific to the US, PS, or CS HIV RNAs as well as a large number of proteins preferentially associated with both the US and PS HIV RNAs, which are Rev-dependent, intron-retaining transcripts responsible for late HIV gene expression (Figure 2A and Table S7). Of the 210 host proteins identified as HIV-1 RNA interactors, 25 had been previously shown to associate with US HIV RNA (Knoener et al., 2017) and 51 with viral proteins (Gag, Gag-Pol, Tat, Rev) that are known to be involved in HIV-1 RNA regulation (Oughtred et al., 2019) (Table S7). Based on siRNA knockdown, at least 48 represent potential new host regulatory factors (Table S7). Using Gene Ontology (GO) term enrichment analysis we showed that several biological processes and cellular components are over-represented in each splice variant subgroup (Figure 2B, Table S4), suggesting cellular pathways that may be uniquely involved in the processing of a subset of HIV splice variants. Notable was enrichment of proteins related to the regulation of mRNA stability and splicing in the US and PS interactomes and proteins related to mitochondrial gene expression and organization in the US interactome.

With a focus on late-stage Rev-dependent US and PS RNA nuclear export, we examined a large cluster of 16 proteins with closely related HIV splice variant interaction profiles which preferentially associated with US and PS, but not CS, transcripts (Figure S7, Table S7). Eleven of these host proteins were known stress granule components and/or had functions in splicing. Ten were determined to be positive effectors of late gene expression, six of which affected late gene expression significantly more than early. Within this group, a subcluster of three proteins (HNRNPR, FAM120A, and IGF2BP3) exhibited markedly similar HyPR-MS RNA interaction profiles (Figure 4C), siRNA KD effects (Figure 4D), and subcellular localization patterns (Figure 4A). HNRNPR and IGF2BP3, as well as IGF2BP1 and YBX1 which also cluster with this group (Figure S7), were previously identified as components of IGF2BP1-ribonucleoprotein granules (IMP1-granules); cytoplasmic granules that contain and confer stability to mRNAs that have not yet been translated (Jonson et al., 2007). HNRNPR was also shown to stabilize and facilitate subcellular localization of RNA (Briese et al., 2018; Reches et al., 2016). Intriguingly, YBX1, IGF2BP1, and HNRNPR have previously been suggested to have roles in HIV replication: YBX1 was shown to stabilize HIV US RNA and enhance virus production (Jung et al., 2018; Mu et al., 2013), overexpression of IGF2BP1 was shown to reduce HIV infectivity through its interaction with Gag (Zhou et al., 2008), and HNRNPR was shown to interact with HIV Rev (Hadian et al., 2009). By contrast, FAM120A has not previously been linked to viruses but has been shown to protect RNAs from Ago2-mediated degradation through the RNA-induced silencing complex (RISC) (Kelly et al., 2019), which frequently serves in an antiviral role (Eckenfelder et al., 2017; Harvey et al., 2011). Our functional analysis showed a decrease in late HIV gene expression upon knockdown of all five of these proteins (IGF2BP1, YBX1, HNRNPR, IGF2BP3, and FAM120A; Table S7), consistent with shared roles for these clustered host proteins as positive regulators of US HIV RNA transport and/or stability.

Analysis of the HyPR-MS and siRNA knockdown screen data also revealed a trend for mitochondria-linked proteins that interacted with US and/or PS HIV RNA and served as negative effectors of HIV gene expression. Of particular interest were LRPPRC and MBOAT7 because they both preferentially interacted with US and PS HIV RNA; were categorized as negative effectors of late gene expression, and could be detected co-localizing with US RNA both in the nucleus and the cytoplasm. LRPPRC was previously shown to localize to the nucleus and to mitochondria as a putative effector of RNA metabolism in both locations (Mili and Pinol-Roma, 2003; Ruzzenente et al., 2012). One study showed that nuclear LRPPRC directly interacted with CRM1, eIF4E, and a signature RNA secondary structure found in a subset of cellular RNAs (Volpon et al., 2017); features similar to how Rev and the RRE are known to drive US and PS RNA export. Interestingly, another study implicated LRPPRC in HIV-1 replication but as affecting the pre-integration stages (Schweitzer et al., 2012). By contast, MBOAT7 is an intramembrane protein and acyltransferase that incorporates polyunsaturated fatty acids into phosphatidylinositol (Lee et al., 2008; Lee et al., 2012); and has not previously been implicated in viral or cellular RNA metabolism. However, in addition to perinuclear and ER localization, MBOAT7 has been reported to localize to mitochondrial associated ER membranes (MAMs) which bridge the ER to the mitochondria to regulate antiviral signaling through the mitochondrial antiviral-signaling (MAVS) viral RNA sensor (Hirata et al., 2013). Based on our combined results, we hypothesize that both LRPPRC and MBOAT7 link US RNA transport to mitochondrial signaling pathways capable of dampening HIV-1 late stage gene expression.

This study describes and validates a powerful new biochemical approach for deep interrogation of the complex interplay of viral and cellular RNA and protein factors during viral infection (Figure 6). Using siRNA KD and single cell imaging experiments we also generated a catalog of host protein candidates for positive and negative regulation of HIV-1 gene expression. Interestingly, a subset of proteins (HNRNPR, RBM4, and RBMX) were frequently observed both at US RNA transcription sites as well as at putative sites of virus particle assembly, suggesting that these factors may be capable of strong, persistent association with viral RNA throughout the entire productive phase (Figure 6A). The changes seen in subcellular distribution of HNRNPR are consistent with a role in stability and nuclear export of intron-retaining HIV-1 transcripts while the expression changes and localization of IGF2BP3 support a role in cytoplasmic US RNA transport and stability (Figure 6B). Finally, we identified a set of host factors linked to mitochondria (including LRPPRC and MBOAT7) that may represent new effectors of HIV-1 antiviral surveillance.

## MATERIALS and METHODS

### Cell Lines

Jurkat cells are T lymphocytes established from the peripheral blood of a 14-year-old male with acute T cell leukemia and were obtained from ATCC (TIB-152). The cells were cultured in RPMI media supplemented with 10% fetal bovine serum and 1% L-glutamine-penicillin-streptomycin in roller bottles rotated at 3 rotations per minute (rpm) at 37°C in 5% CO2. A cell density of 1X10^6^ cells per mL of media was maintained by regular quantification. This cell line was authenticated by karyotyping.

HEK293T cells are human embryonic kidney cells and were obtained from ATCC (CRL-11268). The cells were cultured in DMEM media supplemented with 10% fetal bovine serum and 1% L-glutamine-penicillin-streptomycin at 37°C in 5% CO_2_. This cell line was authenticated by morphology and are G418 resistant.

Human 293T cells stably-expressing YFP-ACT were cultured in Dulbecco’s modified Eagle’s medium (DMEM) supplemented with 10% fetal bovine serum, 1% L-glutamine, and 1% penicillin-streptomycin.

HeLa cells were cultured in DMEM media supplemented with 10% fetal bovine serum and 1% L-glutamine-penicillin-streptomycin at 37°C in 5% CO_2_.

### HIV-1 Virion Production

2.5X10^6^ HEK293T cells were plated in 10 cm tissue-culture treated dishes in 10mL media then transfected using polyethylenimine with 1μg of DNA plasmid expressing the G envelope glycoprotein from vesicular stomatitis virus (VSV-G) and 9μg of plasmid DNA encoding the full-length NL4-3 molecular clone of HIV-1 bearing inactivating mutations in *env, vpr*, and either a.) expressing a Cyan Fluorescent Protein (CFP) reporter from the *nef* reading frame (HIV-1 E-R-CFP) (Adachi et al., 1986; Becker and Sherer, 2017) or b.) expressing mCherry in the nef ORF and three copies of CFP, in tandem, between the matrix and capsid ORFs of Gag (E-R-Gag-3xCFP mCherry/nef) (Hendrix et al., 2015; Holmes et al., 2015; Mergener et al., 1992). At 24-hours post-transfection, media was replaced with 4mL fresh media. At 48-hours post-transfection, culture supernatants were harvested, filtered through a sterile 0.45μm syringe filter and frozen at −80°C. Dose of HIV-1 E-R-CFP viral inoculum required for effective infection was determine in small-scale infection titrations in Jurkat cells.

### HyPR-MS Analysis

#### Jurkat Cell Infections

1X10^8^ Jurkat cells in 25mL RPMI, 25mL viral inoculum (HIV-1 E-R-CFP) in DMEM, and polybrene (concentration 10ug/mL) were combined and incubated in a rotating roller bottle. After three hours, culture volume was increased to 300 mL using RPMI media and incubated at 3 rpm for 45 hours. Infection was confirmed to be >90% by visualizing CFP expression via epifluorescence microscopy. Cells were centrifuged at 1500rpm for 10 minutes, washed three times with PBS, then cross-linked by resuspending in 0.25% formaldehyde and incubated at room temperature for 10 minutes. Cross-linked cells were washed once with PBS then resuspended in 100mM Tris-HCl and incubated at room temperature for 10 minutes to quench formaldehyde. Cells were washed twice more in 1xPBS, pelleted by centrifugation, and frozen at −80°C.

#### Cell Lysis

Jurkat cells pellets were resuspended on ice in lysis buffer (469mM LiCl, 62.5mM Tris HCl, pH 7.5, 1.25% LiDS, 1.25% Triton X-100, 12.5mM Ribonucleoside Vanadyl Complex, 12.5mM DTT, 125U/mL RNasin Plus, 1.25X Halt Protease Inhibitors) to a final cell concentration of 5X10^6^ cells/mL. Cells were lysed by frequent vortexing for 10 minutes, keeping the cells on ice between vortexes.

#### HIV-1 RNA Splice Variant Hybridization and Capture

Each capture replicate used 5 x 10^7^ cells. Three biological replicates of each splice variant capture was conducted for this analysis. The HIV-1 US, PS, and CS RNAs were each purified from the Jurkat cell lysate by three sequential and separate hybridization and capture events; US followed by PS followed by CS HIV RNA. The amounts of biotinylated capture oligonucleotides and streptavidin coated magnetic beads for each hybridization and capture are listed in Table S1. The appropriate concentrations of biotinylated capture oligonucleotides were added to the Jurkat cell lysates and the final concentration of lysis buffer (375mM LiCl, 50mM Tris, 1% LiDS, 1% Triton X-100, 10mM RVC, 10mM DTT, 100U/mL RNasin Plus, 1X Halt Protease Inhibitors) was obtained by addition of nuclease free water. The samples were then incubated at 37°C for three hours with gentle nutation. Streptavidin coated magnetic Speedbeads were washed 3 times with wash buffer (375mM LiCl, 50mM Tris, 0.2% LiDS, 0.2% Triton X-100) prior to addition and nutation for one hour at 37°C with the hybridization samples. Using a magnet, the beads were collected to the side of each tube, and the lysate was removed and transferred to a clean tube for the next hybridization and capture. The beads were then washed 2 times, for 15 minutes each, at 37°C with a volume of wash buffer 5-times the volume of the original aliquot of beads used for capture (i.e. 5X bead volume) then one time for 5 minutes at room temperature with a 5X bead volume of release buffer (100mM LiCl, 50mM Tris, 0.1% LiDS, 0.1% Triton X-100).

#### Release of HIV RNA from Beads

The beads for the US, PS, and CS RNA captures were individually resuspended in a 3X bead volume of release buffer. The appropriate amount of release oligonucleotide (Table S1) was added and the bead mixture was nutated at room temperature for 30 minutes. Using a magnet to collect the beads to the side of the tube, the supernatant containing the released RNA-protein complexes was transferred to a clean tube. The resulting sample was divided into two aliquots; 2% for RT-qPCR analysis and 98% for mass spectrometric analysis.

#### RNA Extraction and Reverse Transcription

2% by volume of each release sample was incubated overnight at 37°C with 1 mg/mL proteinase K, 4mM CaCl2, and 0.2% LiDS to remove the proteins. The RNA was then extracted from the samples using TriReagent per manufacturer’s protocol and precipitated in 75% ethanol, with 2uL of GlycoBlue, at −20°C for at least 2 hours. The RNA was pelleted by centrifugation at 20,800 g and 4°C for 15 minutes, the pellet was washed with 75% ethanol, centrifuged at 20,800 g and 20°C for 15 minutes, then resuspended in 15uL of nuclease free water. 10uL of the purified RNA was used for reverse transcription (High Capacity cDNA Reverse Transcription Kit, Applied Biosystems) per the manufacturer’s protocol. The procedures described here were also performed on HIV-1 E-R-CFP virus inoculum for isolation and analysis of a semi-purified standard of the US HIV RNA. The isolated RNA was quantified by NanoDrop analysis, serially diluted, reverse transcribed and then used for a standard calibration curve for qPCR analysis.

#### qPCR Analysis

The 20uL reverse transcription product was diluted with 20uL of nuclease free water and analyzed using sequence-specific qPCR primers and probes (Table S1) and Roche LightCycler 480 Probes Master Mix for relative quantitation of the US, PS, and CS HIV transcripts and human GAPDH. Purified HIV-1 E-R-CFP plasmid was quantified by NanoDrop analysis, serially diluted, then used as a standard calibration curve for qPCR analysis.

#### Protein Purification and Trypsin Digestion

98% by volume of each capture sample was processed using an adapted version of eFASP (Erde et al., 2014) for purification of proteins. Amicon 50kDa MWCO filters and collection tubes were passivated by incubating overnight in 1% CHAPS and then rinsed thoroughly with mass spectrometry grade water. Each release sample was brought to a final concentration of 8M Urea and 0.1% deoxycholic acid (DCA) then passed through the filter in 500uL increments by centrifugation for 10 minutes at 14,000 g. RNA-protein complexes were trapped in the filter and the eluent passed through to a collection tube for discarding. In the same manner (addition of solution followed by centrifugation), the following passages were conducted: 1.) Three passages of 400uL of exchange buffer (8M urea, 0.1% DCA, 50mM Tris pH 7.5), 2.) Incubation for 30 minutes with 200uL of reducing buffer (8M urea, 20mM DTT), 3.) Incubation for 30 minutes, in the dark, with alkylation buffer (8M urea, 50mM iodoacetamide, 50mM ammonium bicarbonate), 4.) Three passages of 400uL of digestion buffer (1M urea, 50mM ammonium bicarbonate, 0.1% DCA). Finally, the sample remaining in the filter was brought to 100uL with digestion buffer, the filter was transferred to a clean, passivated collection tube and 1ug of trypsin added to the filter for digestion. The filter-collection tube containing the sample was sealed with parafilm to prevent evaporation during incubation overnight at 37°C. Following digestion, the filter-collection tube was centrifuged for 10 minutes at 14,000 g. 50uL of 50mM ammonium bicarbonate was added to the filter followed by centrifugation at 14,000 g for 10 minutes. This step was repeated once to ensure the collection of the entire peptide sample. The 200uL peptide sample was then brought to 1% TFA followed by addition of 200uL of ethyl acetate. The sample was vortexed for 1 minute then centrifuged at 15,800 g for 2 minutes. The top layer was aspirated and discarded and extraction with 200uL ethyl acetate was repeated 2 times. The aqueous layer was then dried using a Savant SVC-100H SpeedVac Concentrator and the sample resuspended in 150uL 0.1% TFA. For removal of salts from the sample a C18 solid-phase extraction pipette tip was first conditioned with 70% ACN, 0.1% TFA, and then equilibrated with 0.1% TFA. The peptide sample was then loaded onto the C18 solid phase by repeated passing of the 150uL sample over the cartridge. The C18 extraction pipette tip was then rinsed with 0.1% TFA 10 times followed by peptide elution in 150μL 70% ACN, 0.1% TFA. The samples were then dried using the SpeedVac Concentrator and reconstituted in 95:5 H2O:ACN, 0.1% formic acid.

#### Mass Spectrometry of Peptides

The samples were analyzed using an HPLC-ESI-MS/MS system consisting of a high performance liquid chromatograph (nanoAcquity, Waters) set in line with an electrospray ionization (ESI) Orbitrap mass spectrometer (LTQ Velos, ThermoFisher Scientific). A 100 μm id X 365 μm od fused silica capillary micro-column packed with 20 cm of 1.7 μm-diameter, 130 Angstrom pore size, C18 beads (Waters BEH) and an emitter tip pulled to approximately 1 μm using a laser puller (Sutter Instruments) was used for HPLC separation of peptides. Peptides were loaded on-column with 2% acetonitrile in 0.1% formic acid at a flow-rate of 400nL/minute for 30 minutes. Peptides were then eluted at a flow-rate of 300 nL/minute over 120 min with a gradient from 2% to 30% acetonitrile, in 0.1% formic acid. Full-mass profile scans were performed in the FT orbitrap between 375-1500 m/z at a resolution of 120,000, followed by MS/MS HCD scans of the ten highest intensity parent ions at 30% relative collision energy and 15,000 resolution, with a mass range starting at 100 m/z. Dynamic exclusion was enabled with a repeat count of one over a duration of 30 seconds. The Orbitrap raw files were analyzed using MaxQuant (version 1.5.3.30) (Cox and Mann, 2008) and searched with Andromeda (Cox et al., 2011) using the combined Uniprot (Breuza et al., 2016) canonical protein databases for human and HIV-1 and supplemented with common contaminants (downloaded June 8, 2016). Samples were searched allowing for a fragment ion mass tolerance of 20 ppm and cysteine carbamidomethylation (static) and methionine oxidation (variable). A 1% false discovery rate for both peptides and proteins was applied. Up to two missed cleavages per peptide were allowed and at least two peptides were required for protein identification and quantitation. Protein quantitation was achieved using the sum of the peptide peak intensities for each protein of each biological replicate and capture sample type. The peak intensities of HIV capture samples were normalized by the total peak intensity of all HIV capture samples and the same was done for scrambled capture samples.

#### MS Data Analysis

To determine the differential interactomes of the HIV-1 splice variants pairwise comparisons (US vs PS, US vs CS, PS vs CS) were statistically analyzed with the student’s T-test and a permutation based FDR (5% threshold) using Perseus software (Tyanova et al., 2016) (Table S3). Proteins that met this threshold in at least one pairwise comparison were then hierarchically clustered using Cluster software (de Hoon et al., 2004) and TreeView (Saldanha, 2004) was used to facilitate visualization of the differential interactomes. Gene Ontology analysis, using PANTHER (Mi et al., 2017), of the proteins statistically elevated in each individual splice variant capture were evaluated for enrichment of terms in the categories of Biological Processes and Cellular Component (Table S4)

### siRNA Knockdown Screen

#### Cell Culture, KD, and Infection

The virus used for determining the effect of gene specific siRNA knockdown on early and late HIV-1 gene expression was a two-color fluorescent HIV-1 reporter virus (E-R-Gag-3xCFP mCherry/nef). This virus expresses mCherry in the nef ORF and three copies of CFP, in tandem, between the matrix and capsid ORFs of Gag, in a similar but expanded manner as previously done (Hendrix et al., 2015; Holmes et al., 2015; Mergener et al., 1992). This virus allows screening for early (mCherry; completely-spliced gene products) and late (CFP; unspliced gene products). Stocks of viral inoculum were produced in 293T cells by transfecting the E-R-Gag-3xCFP mCherry/nef with psPAX2 and VSV-G. Human 293T cells stably-expressing YFP-ACT were cultured in DMEM supplemented with 10% fetal bovine serum, 1% L-glutamine, and 1% penicillin-streptomycin. All cell incubations during the siRNA KD process were done at 37°C and 5% CO2 in a humidified incubator. Approximately 5×10^3^ cells were plated in 24 wells of a 96-well culture plate and incubated 24 hours; then the media was replaced with 125uL of anti-biotic free DMEM. 0.875uL of DarmaFECT transfection reagent in 25uL of Opti-MEM was mixed with 25uL of Opti-MEM containing 4.4 pmol of gene specific siRNA (Table S5), incubated at room temperature for 20 minutes, then added to the appropriate well of the 96-well plate. The final, in-well concentration of each siRNA was 25nM. After four hours of incubation, the media was replaced with fresh DMEM media and the cells were incubated overnight. 24-hours post transfection the cells were lifted from the bottom of the well by gentle pipetting, divided equally into two wells, incubated for another 24-hours, then again each well containing cells was divided equally into two wells (now a total of four wells for each gene specific siRNA KD). The cells were allowed to adhere for 2-4 hours then a second siRNA transfection as described above was conducted in all four wells. Four hours post transfection the siRNA containing media was replaced with fresh media. Additionally, in two of the four wells, polybrene was added to the media (final concentration of 2ug/mL) followed by the HIV-1 reporter virus (E-R-Gag-3xCFP mCherry/nef) inoculum in DMEM. After 24 hours the media was exchanged for fresh media and 48-hours post infection the cells were washed with PBS and fixed for 12 minutes using 4% paraformaldehyde (PFA) in PBS then stored at 4C in PBS until imaged. Two biological replicates, each consisting of two technical replicates of infected and two technical replicates of uninfected cells, were obtained for each siRNA targeted gene. Biological replicates are defined as full siRNA knockdown procedures, from cell plating to cell fixation, performed on different days.

#### Imaging

Imaging experiments were performed on a Nikon Ti-Eclipse inverted wide-field epifluorescence deconvolution microscope (Nikon Corporation). Images were collected using an Orca-Flash 4.0 C11440 (Hamamatsu Photonics) camera and Nikon NIS Elements software (v 4.20.03) using Nikon 4x/0.13 (Plan Apo) objective lense and the following excitation/emission filter set ranges (wavelengths in nanometers): 418 to 442/458 to 482 (CFP), 490 to 510/520 to 550 (YFP), 555 to 589/602 to 662 (mCherry).

#### Image Processing and HIV Gene Expression Quantitation

Images were processed and analyzed using FIJI/ImageJ2 (Rueden et al., 2017). For each well, only cell monolayers were used for quantitation of fluorescence. Cell viability for each gene specific siRNA knockdown was assessed using the ACT-YFP marker. Wells that had YFP fluorescence detected within +/- 1.5 standard deviations of the plate mean were considered acceptable for further analysis. CFP and mCherry fluorescence for each well was normalized to the YFP fluorescence for the gene specific siRNA and negative control siRNA (included in each 96-well plate). The Student’s t-test calculation was performed to determine if a statistically significant change in CFP or mCherry expression was detected between each gene specific siRNA KD and the negative control siRNA KD (Table S6; p-value < 0.05).

#### Western Blot Validation

293T cells, with and without gene specific siRNA knockdown, were lysed in 1x radioimmunoprecipitation assay (RIPA) buffer (10 mM Tris-HCl [pH 7.5], 150 mM NaCl, 1 mM EDTA, 0.1% SDS, 1% Triton X-100, 1% sodium deoxycholate) and sonicated. Samples were then boiled for 10 minutes in 2x dissociation buffer (62.5 mM Tris-HCl [pH 6.8], 10% glycerol, 2% sodium dodecyl sulfate [SDS], 10% β-mercaptoethanol), run on SDS-PAGE 10% polyacrylamide gels, and transfered to nitrocellulose membranes (0.2 μM pore size). Immunoblotting was performed as previously described (Becker and Sherer, 2017; Behrens et al., 2017; Garcia-Miranda et al., 2016) using the primary and secondary antibodies detailed in Table S8.

### Co-localization and Expression Quantitation

#### Cell Culture and Infection

HeLa cells cultured in DMEM in 8-well Ibidi plates were infected with HIV-1 attenuated virus (E-R-CFP), described above, by adding polybrene to each well at a final concentration of 2ug/mL followed by the virus inoculum. Media was exchanged with fresh media 24-hours post-infection (h.p.i). The cells were fixed 48-hours h.p.i by washing with PBS, incubating with 3.7% formaldehyde for 10 minutes at room temperature, then washing three times with PBS. Cells were then made permeable by incubating with 0.2% Triton X-100 for 15 minutes at room temperature and washing three times with PBS. Endogenous RNases were then deactivated by incubating with 0.1% DEPC in PBS for 15 minutes, removing the solution then incubating again with fresh 0.1% DEPC in PBS for 15 minutes; the cells were then washed three times with PBS and stored at 4°C.

#### Immunofluorescence (IF) Labeling

All immunofluorescence steps were conducted at room temperature. Blocking buffer was added to each well, incubated for 30-60 minutes, then removed. Cells were then incubated with fresh blocking buffer containing the appropriate primary antibodies at designated concentrations (Table S8) for 60 minutes followed by four, 5 minute, washes with blocking buffer. Blocking buffer containing appropriate concentrations of the secondary antibodies and DAPI stain (Table S8) were then incubated with the cells for 40 minutes followed by 4, 5 minute, washes with PBS. Finally, the cells were fixed with 3.7% formaldehyde for 10 minutes followed by 3 washes with PBS.

#### Fluorescence In Situ Hybridization (FISH)

The FISH protocol was conducted using Stellaris designed hybridization probes (Table S10) and Stellaris FISH reagents. All FISH steps were conducted in the dark. Cells were washed with FISH Wash Buffer A for 5 minutes at room temperature. FISH Hybridization Buffer containing 12.5nM FISH probes was added to each well and incubated at 37°C for 4 hours. The cells were then washed twice with FISH Wash Buffer A for 30 minutes at 37°C then washed once with FISH Wash Buffer B for 5 minutes at room temperature.

#### Order of Protocols

The performance of each primary antibody was dependent on the order that the FISH and IF protocols were performed. For some protein/antibody pairs (DNM2, HNRNPR, FAM120A, MBOAT7, MOV10, RBM4, RBMX) the IF signal was superior if the IF was conducted prior to FISH. For other antibodies (CSDE1, LRPPRC, TRIM56) the IF signal was superior if the IF was conducted after FISH. For G3BP1 and IGF2BP3, either order was fine. The protocols for each procedure remained consistent, the order in which they were done was only reversed.

#### HIV RNA, Gag, and Host Protein Single-Cell Imaging

Single-cell imaging experiments were performed on a Nikon Ti-Eclipse inverted wide-field epifluorescence deconvolution microscope (Nikon Corporation). Images were collected using an Orca-Flash 4.0 C11440 (Hamamatsu Photonics) camera and Nikon NIS Elements software (v 4.20.03) using Nikon 60x (N.A. 1.40; Plan Apo) or 100X (N.A. 1.45; Plan Apo) objective lenses and the following excitation/emission filter set ranges (wavelengths in nanometers): 405/470 (DAPI), 430/470 (CFP), 490/525 (AlexaFluor488), 585/610 (CAL Fluor Red 590), 645/705 (AlexaFluor647). Images were generally acquired in z-stacks containing various numbers of images along the z-axis of the cells. Images were processed and analyzed using FIJI/ImageJ2 (Rueden et al., 2017). All z-frames within a z-stack were examined for instances of co-localization; however, the fluorescence from only a single z-frame was used to produce co-localization images. For determining HIV RNA, Gag, and host protein expression differences in cells, four z-frames were merged additively for fluorescence quantitation of each component.

#### Quantitation of HIV RNA, Gag, and Host Protein Immunofluorescence

Fluorescence for each channel (HIV-RNA, Gag protein, and each host protein) was quantified using FIJI/ImageJ2 (Rueden et al., 2017). Nuclear and total cellular fluorescence were measured by drawing perimeters around the nucleus (defined by DAPI staining) and the entire cell (defined by Gag protein fluorescence in late stage cells or autofluorescence in uninfected/early stage cells) then using FIJI quantitation tools to measure the fluorescence within each drawn perimeter. Cytoplasmic fluorescence was calculated by subtracting nuclear fluorescence from total cell fluorescence and the Nuc/Cyto ratio was calculated by dividing the nuclear fluorescence by the cytoplasmic fluorescence (Table S13). For determining correlation of host protein expression with HIV gRNA expression, cells with outlier values in total HIV gRNA fluorescence were excluded from the dataset. An outlier here is defined as a value that is more than 1.5 interquartile ranges (IQRs) below the 1^st^ quartile (Q1) or above the 3^rd^ quartile (Q3). IQR is defined as (Q3 – Q1); with Q3 and Q1 calculated using the quartile function in Excel (Table S13). The Pearson’s R value was calculated, excluding outliers, to determine correlation of US HIV RNA and host protein fluorescence expression in the nucleus, cytoplasm, total cell, and for the Nuc/Cyto ratios using the CORREL function in Excel. R^2^ values were calculated using the chart tools in Excel (Table S13).

For determining host protein expression and distribution changes outliers were determined, as described above, for host protein expression values in Early and Late cells in four categories: Nuclear, Cytoplasmic, Total Cellular, and Nuc/Cyto ratio. To determine if “Early” and “Late” cells showed statistically significant differences in host protein expression a student’s T-test, excluding outliers, was used to determine a p-value. The percent change in each category was calculated using the mean values for each category within Early and Late cells.

## QUANTIFICATION AND STATISTICAL ANALYSIS

Statistical methods are described in the appropriate “Method Details” section or figure captions for all data analyses.

## ACKNOWLEDGEMENTS

This study was supported by National Institutes of Health [R01AI110221, U54AI150470 to N.M.S., R01CA193481 to L.M.S., T32CA009135 to E.L.E.III]; the Greater Milwaukee Foundation’s Shaw Scientist Program [to N.M.S.]; a UW-Madison UW2020 Infrastructure Award [to N.M.S]; two National Science Foundation Graduate Research Fellowships [DGE-1256259 to J.T.B. and B.E.B.], an OVCGRE Dissertation Completion Fellowship [to J.T.B.]; an Advance Opportunity Fellowship from the UW-Madison SciMed/GRS program [to E.L.E.III].; Any opinions, findings, and conclusions or recommendations expressed in this material are those of the authors and do not necessarily reflect the views of the National Science Foundation.

## AUTHOR CONTRIBUTIONS

Conceptualization, N.M.S., L.M.S., and R.A.K.; Methodology, N.M.S., L.M.S., and R.A.K.; Formal Analysis, R.A.K.; Investigation, R.A.K., E.L.E., J.T.B., M.S., and B.E.B.; Writing-Original Draft, R.A.K.; Writing-Review & Editing, L.M.S., N.M.S., R.A.K., E.L.E., J.T.B., and B.E.B.; Visualization, R.A.K.; Supervision, L.M.S. and N.M.S.; Funding Acquisition, L.M.S. and N.M.S.

## DECLARATION of INTERESTS

The authors declare no competing interests.

---

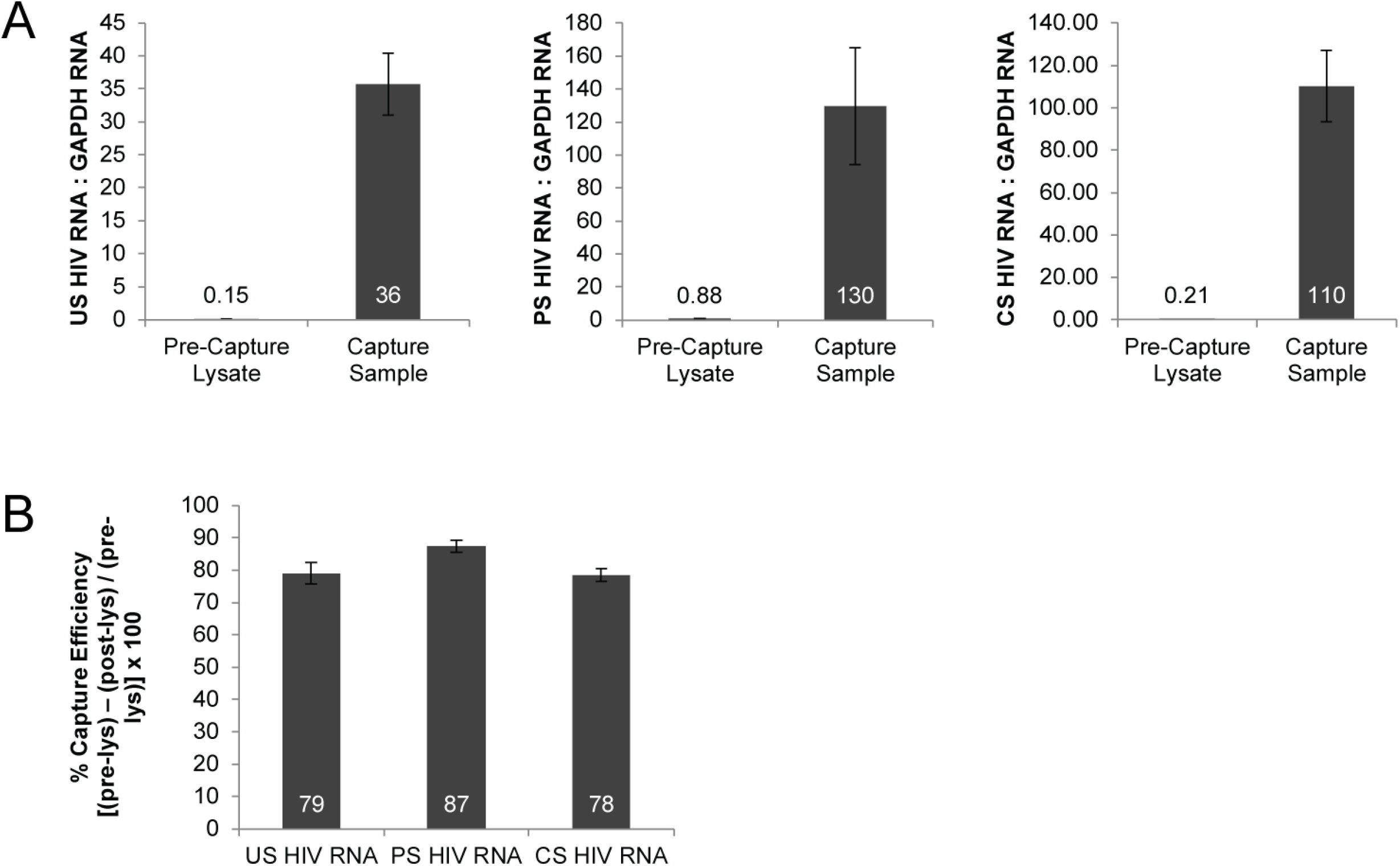

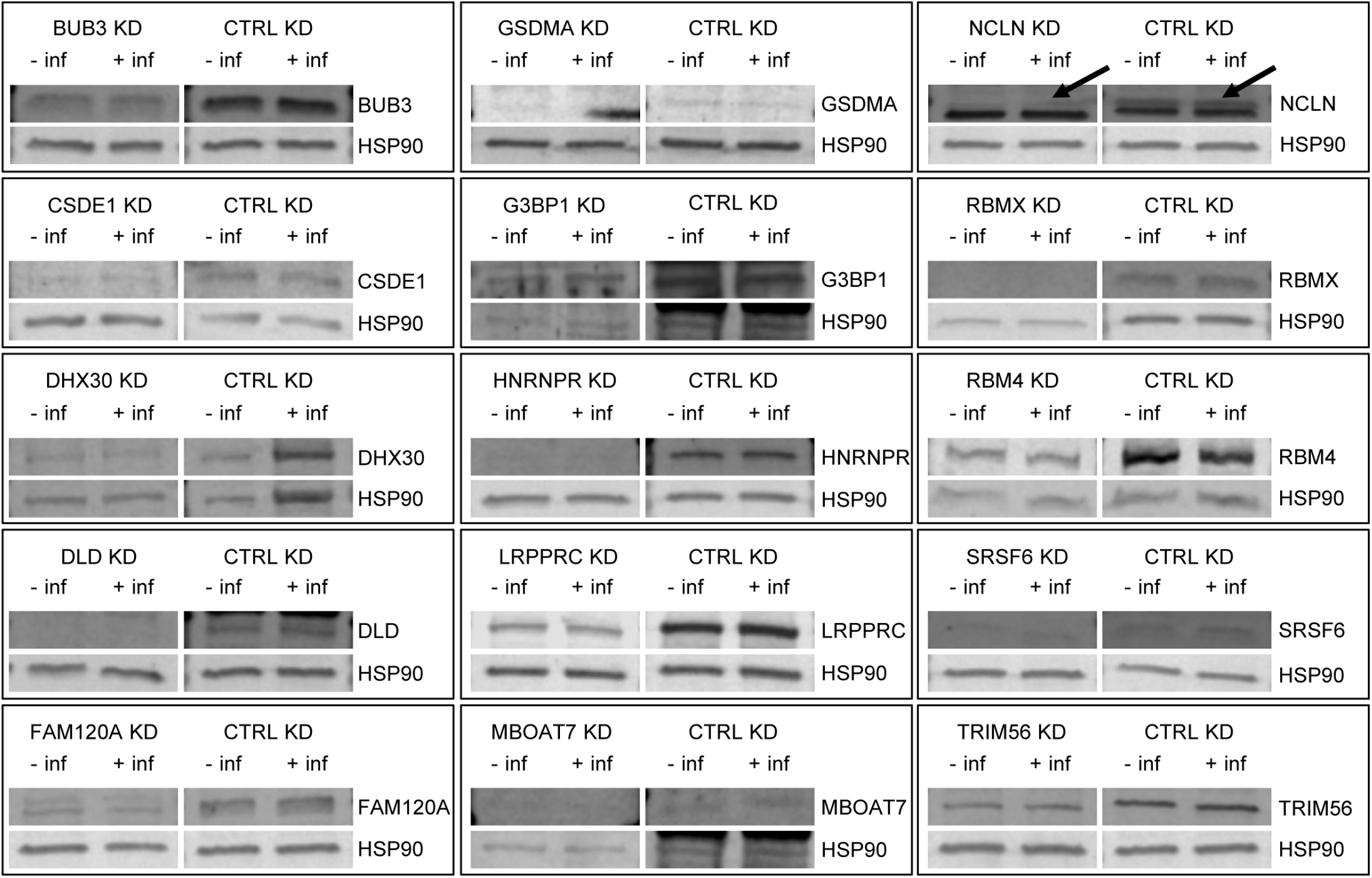

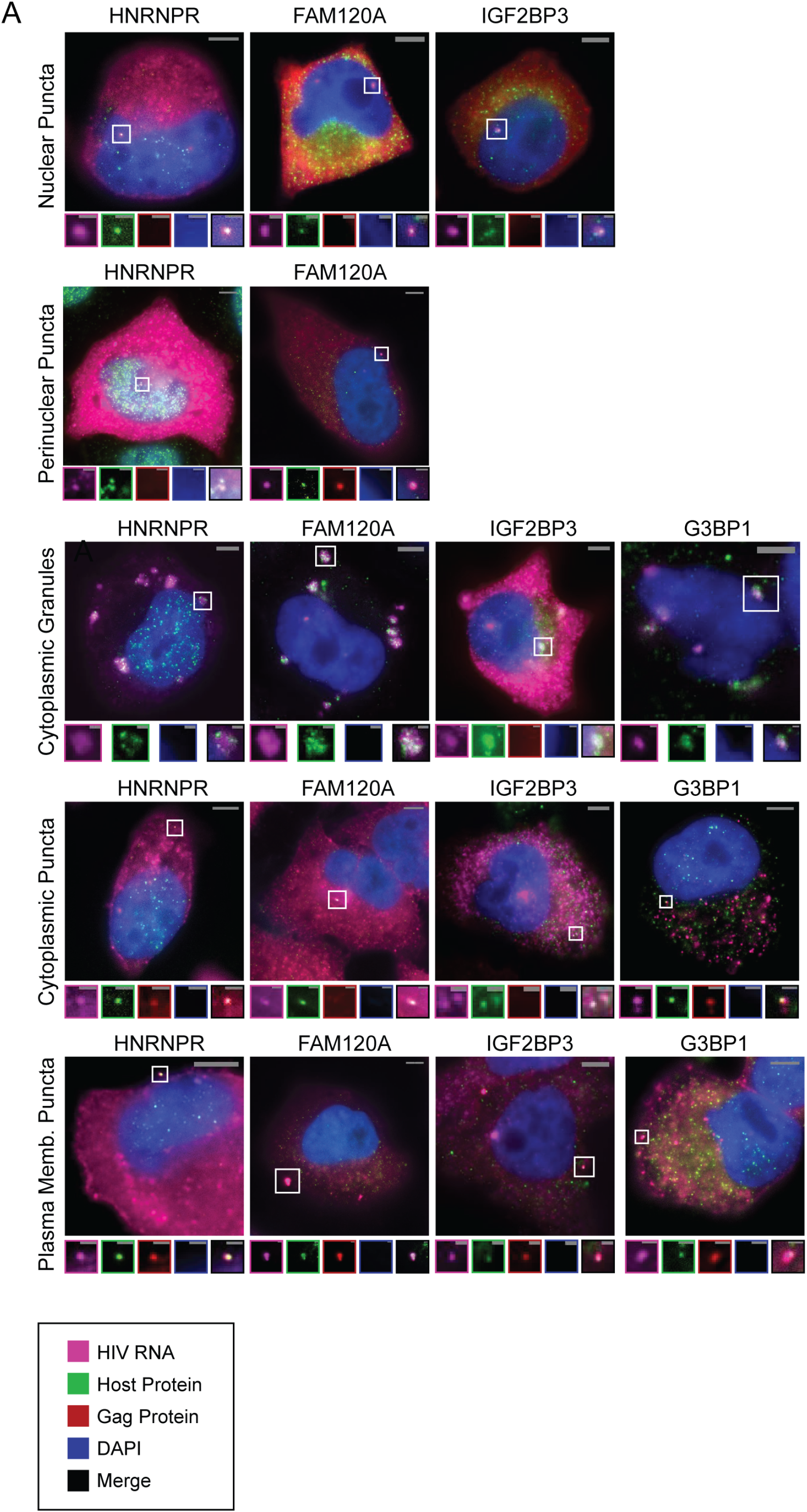
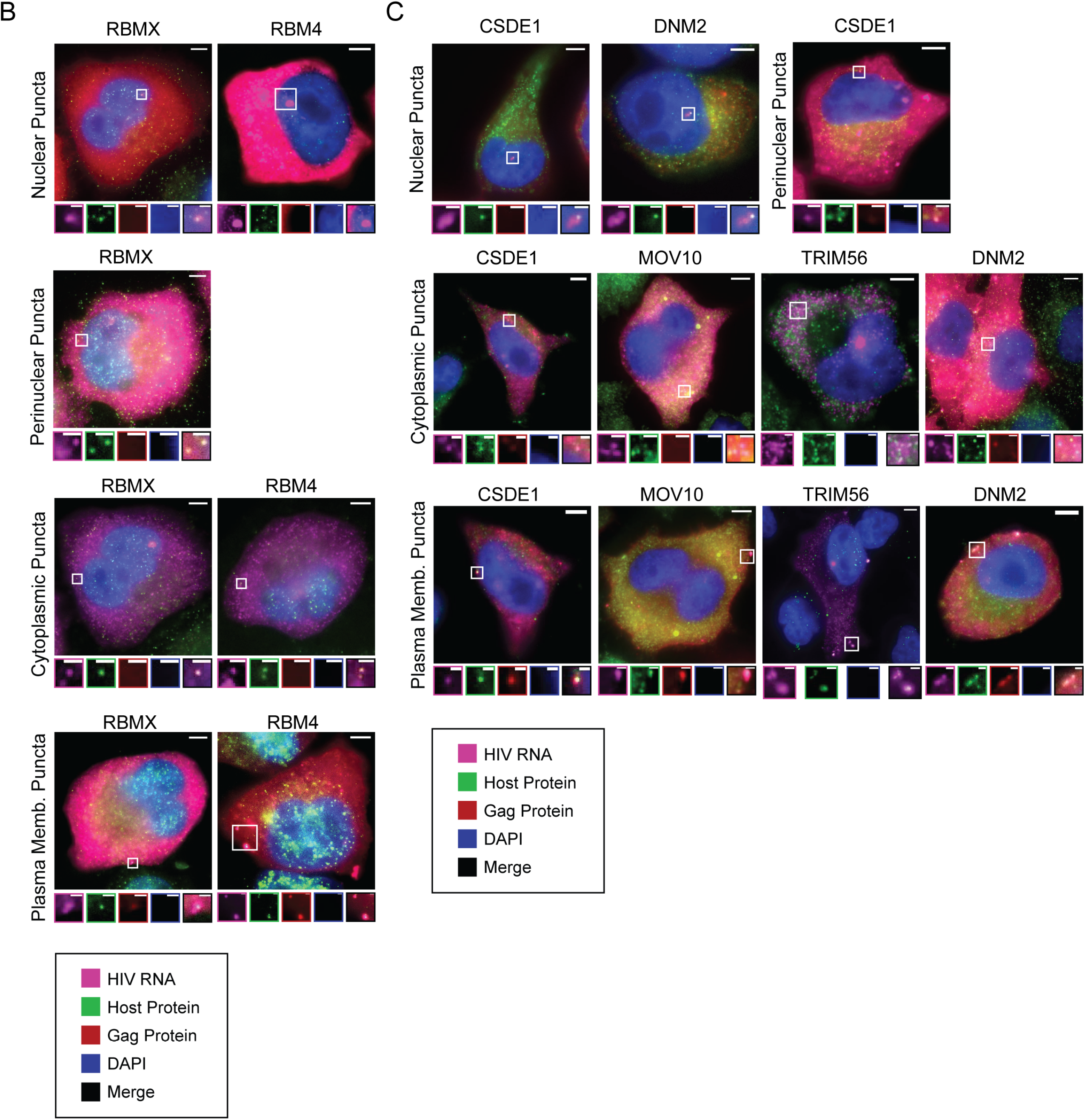
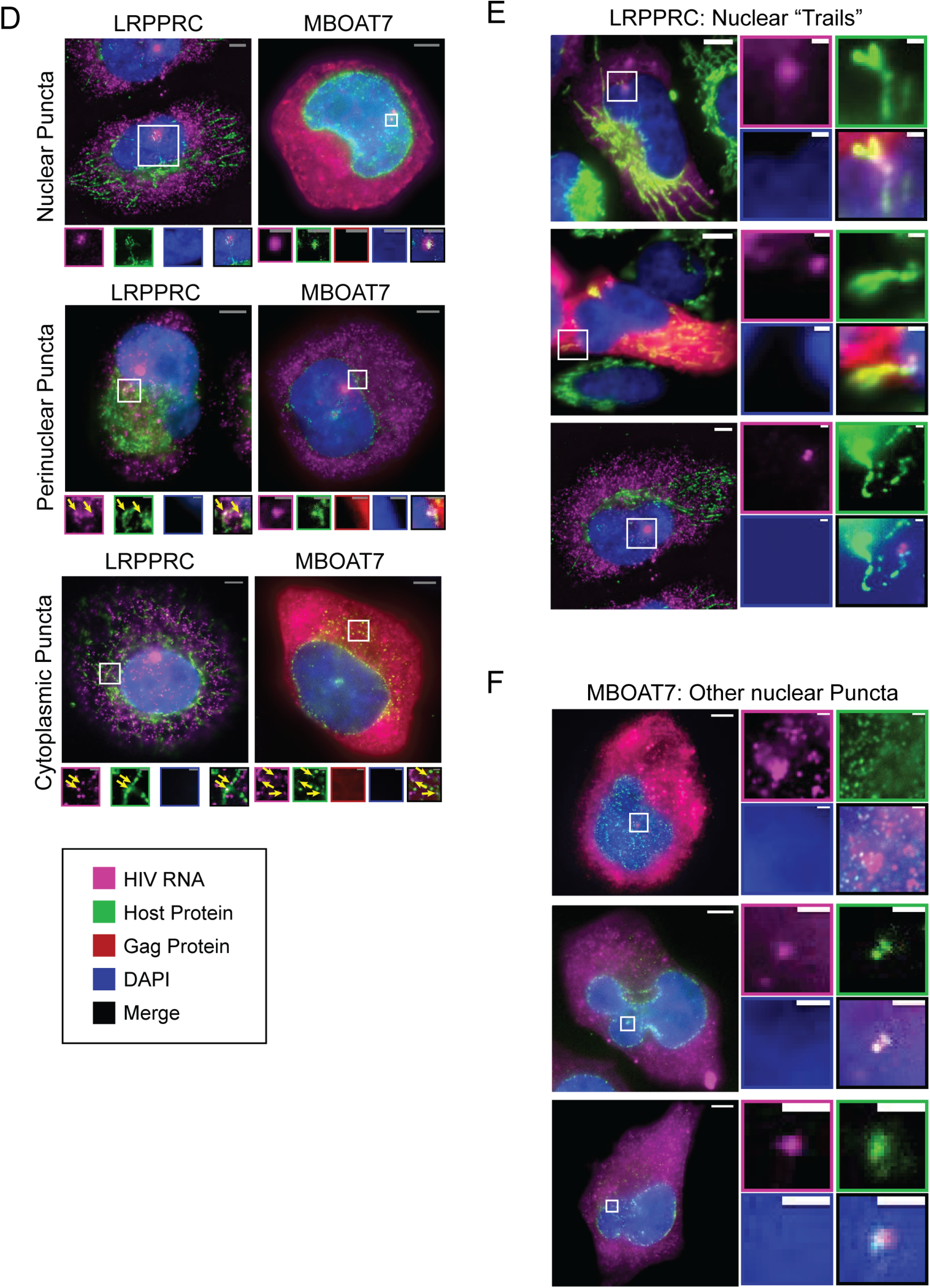

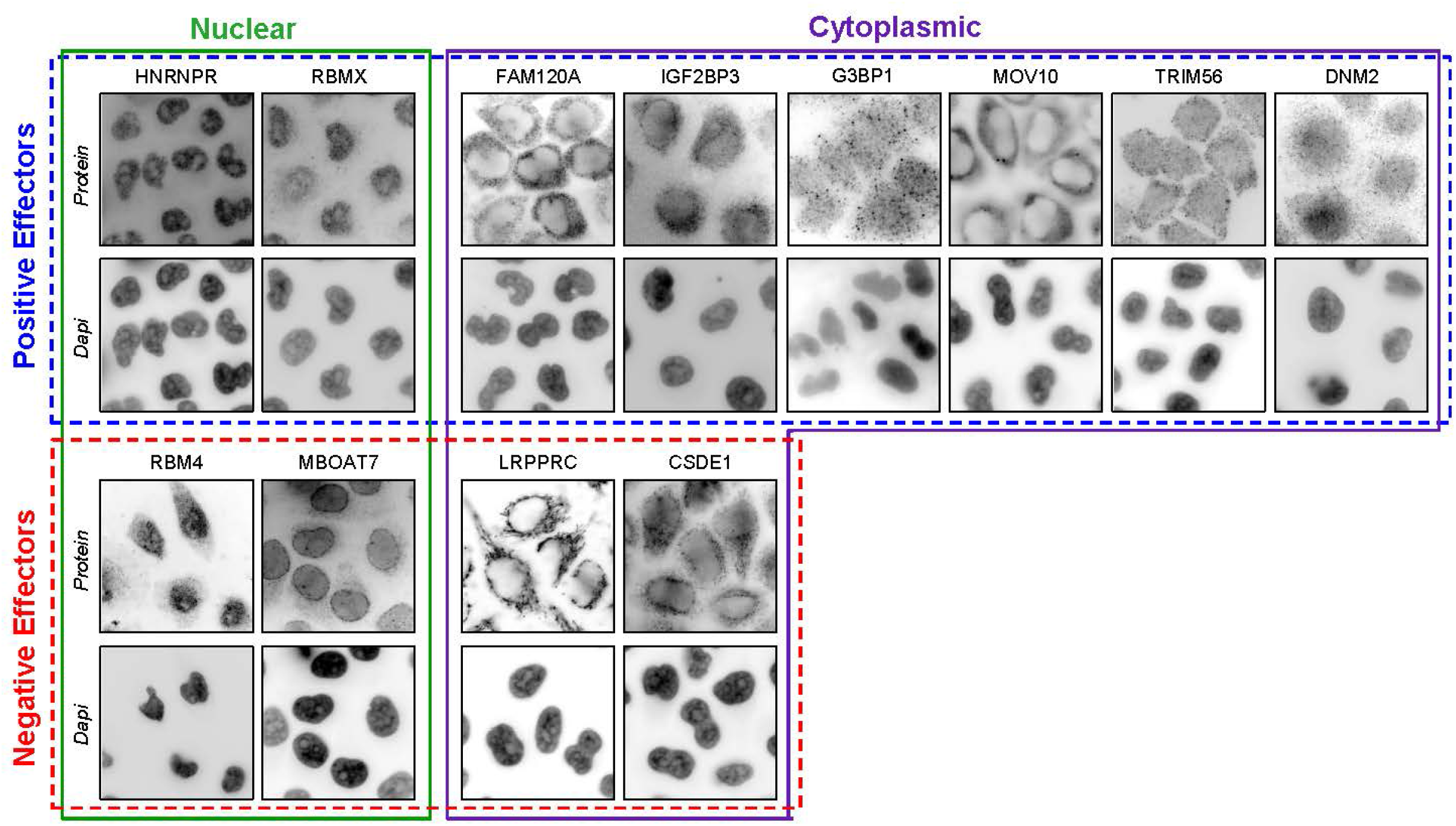

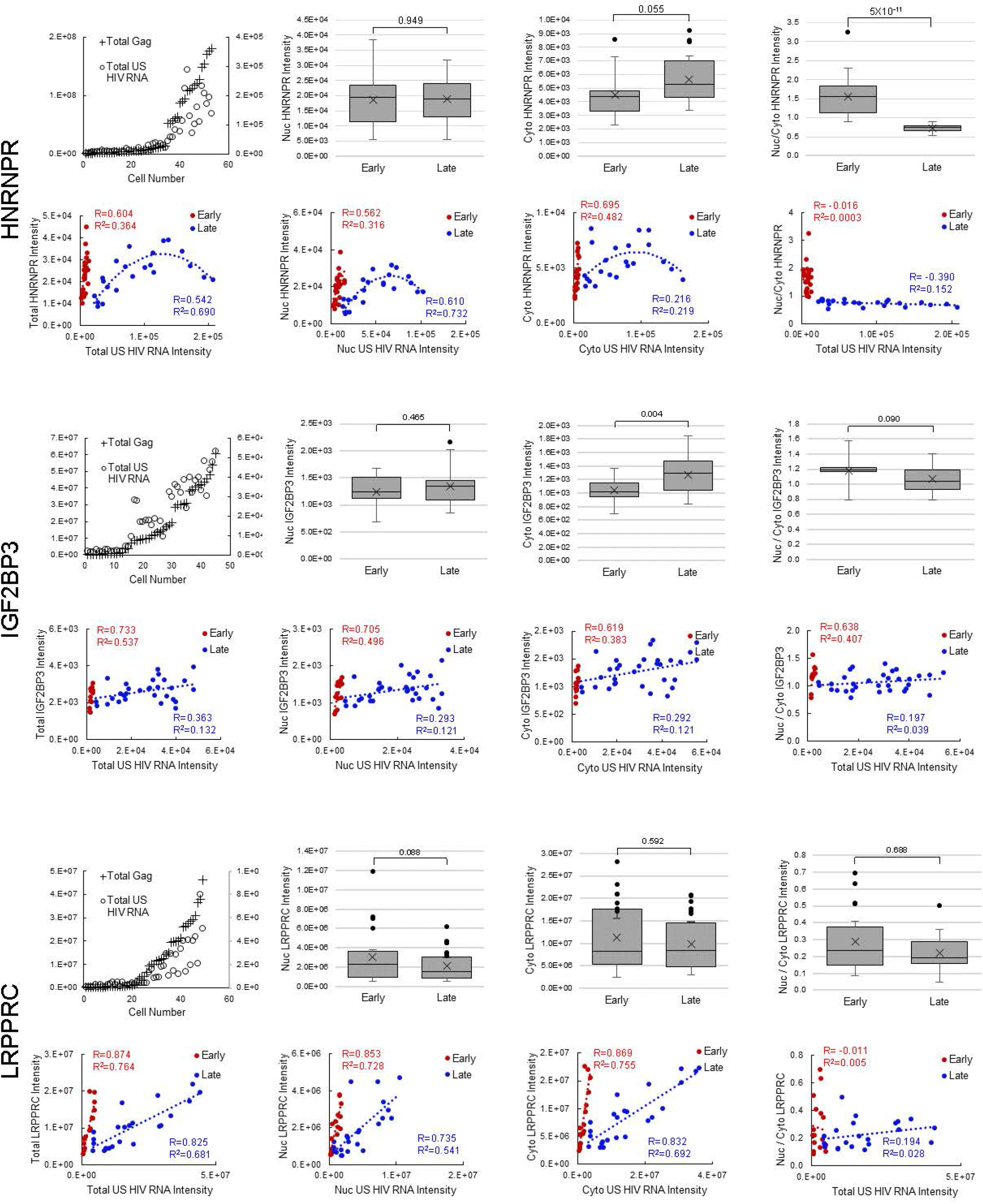

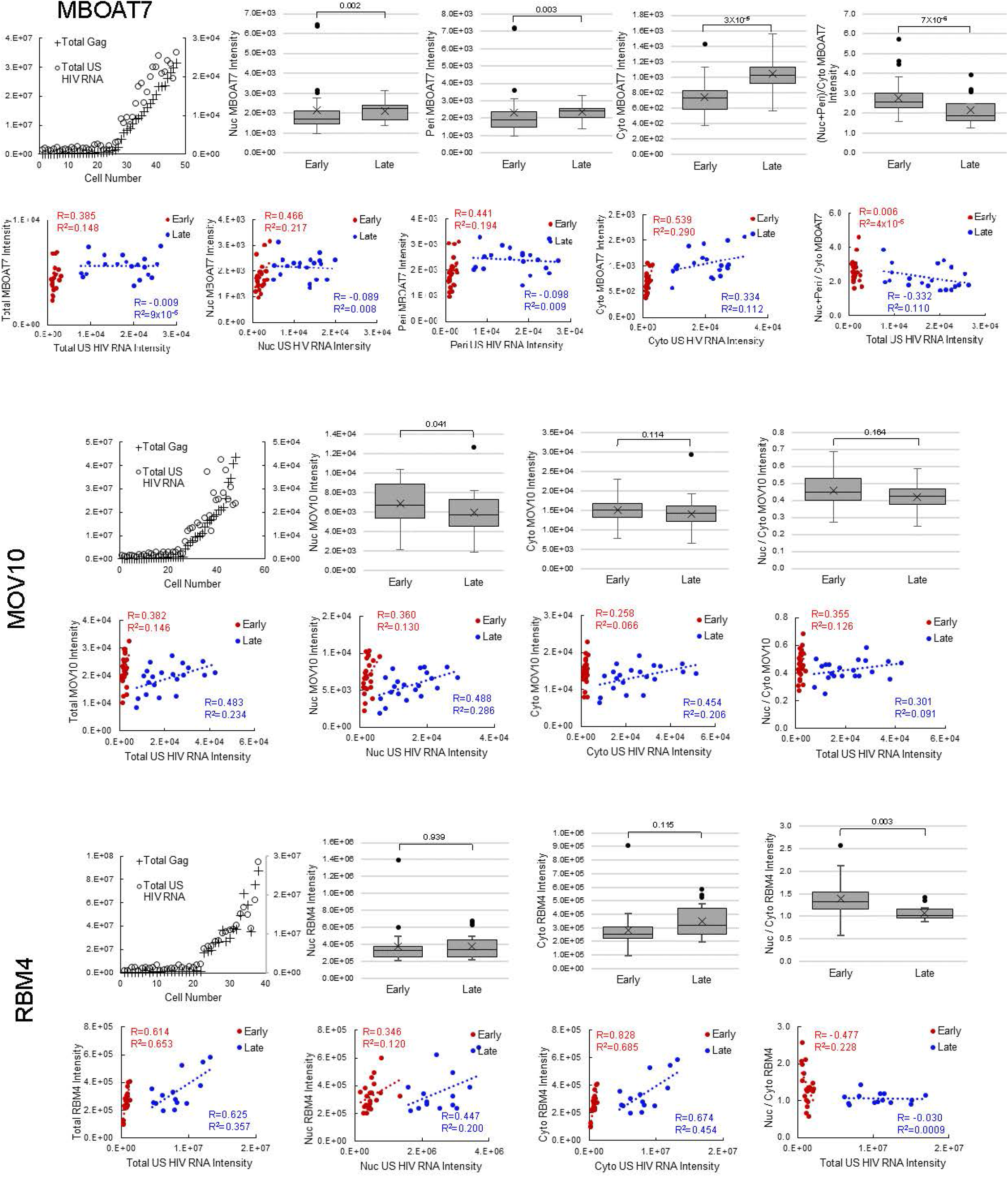

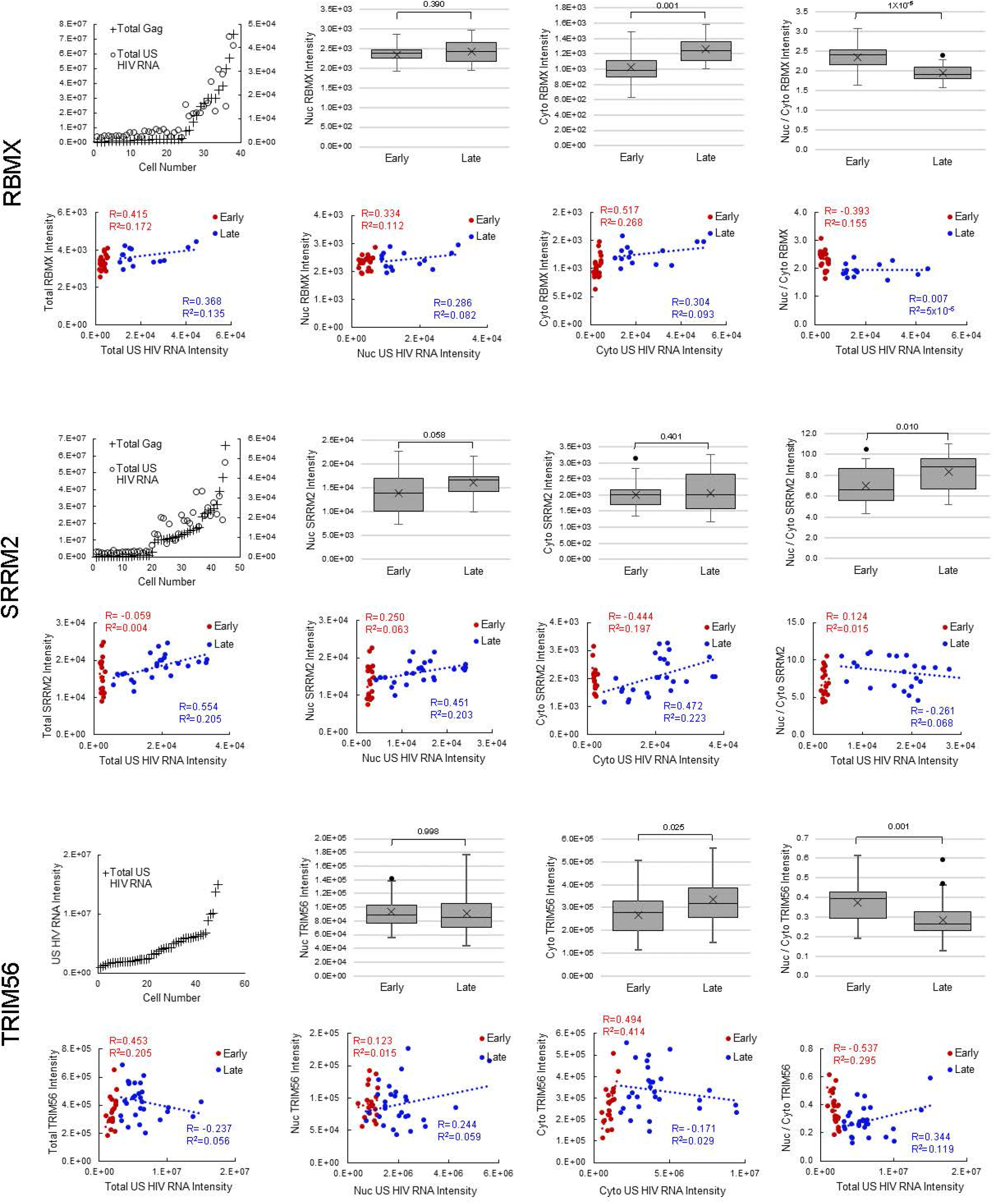

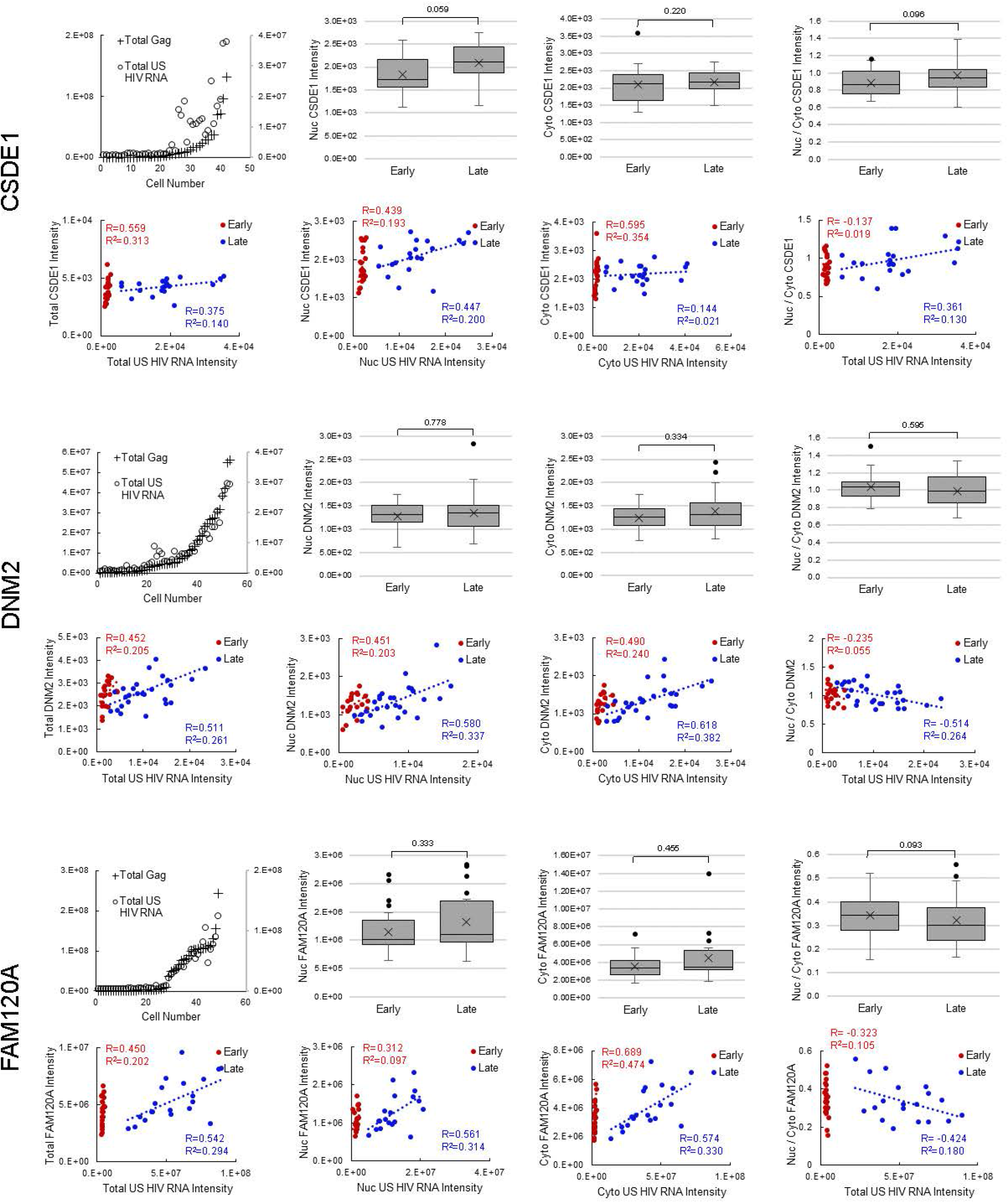

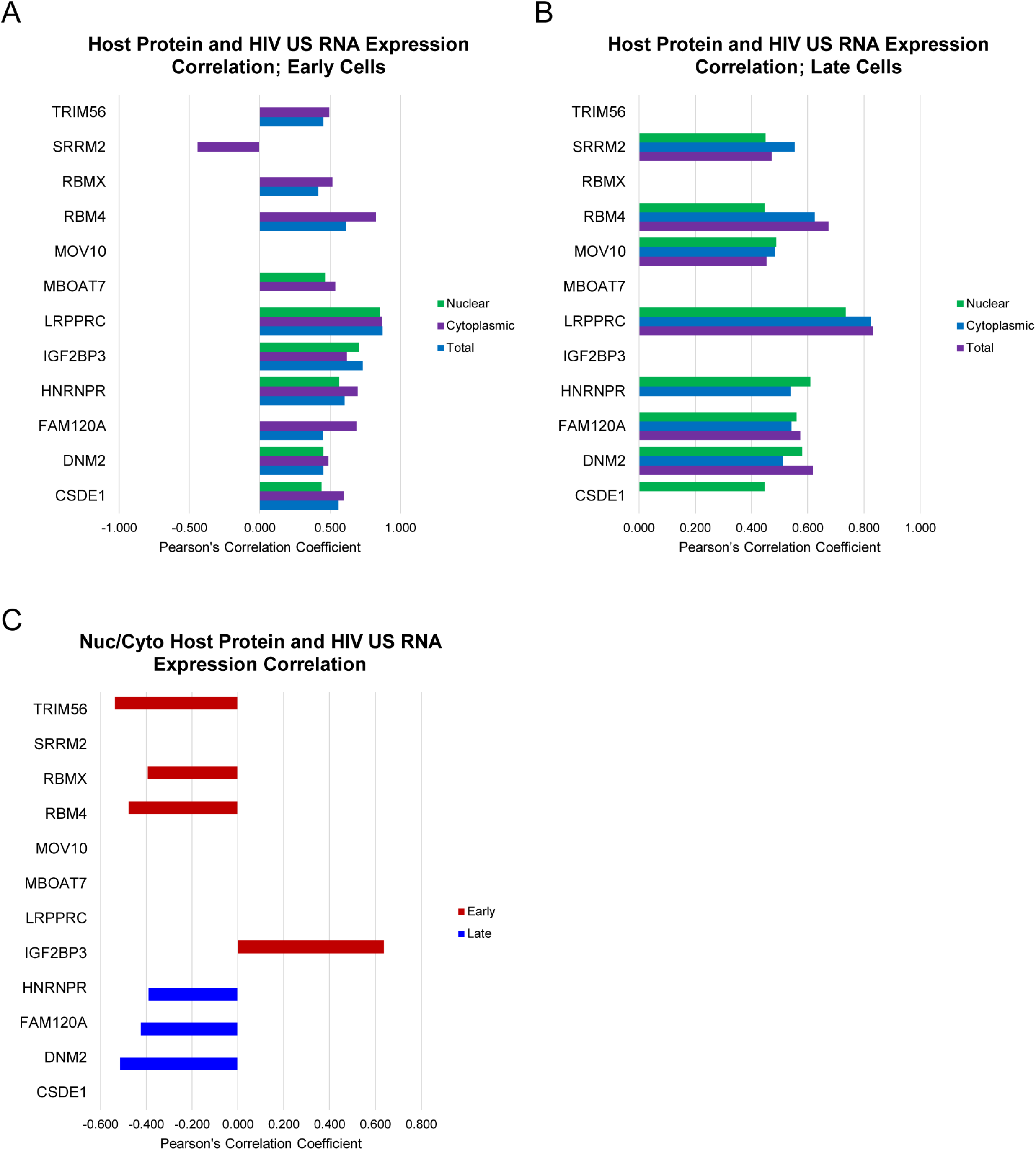

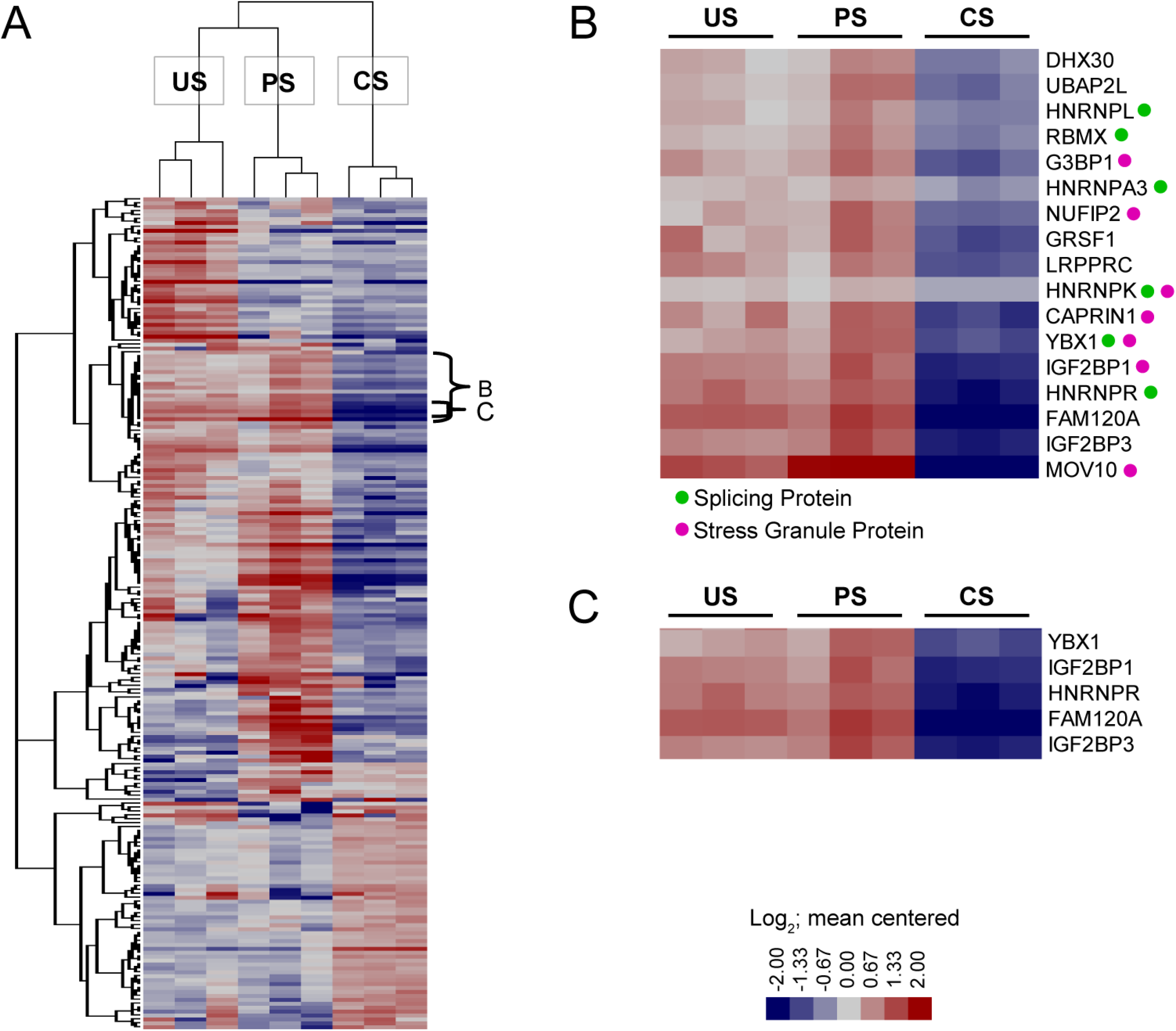

